# Immuno-informatics Design of a Multimeric Epitope Peptide Based Vaccine Targeting SARS-CoV-2 Spike Glycoprotein

**DOI:** 10.1101/2020.07.30.228221

**Authors:** Onyeka S. Chukwudozie, Clive M. Gray, Tawakalt A. Fagbayi, Rebecca C. Chukwuanukwu, Victor O. Oyebanji, Taiwo T. Bankole, A. Richard Adewole, Eze M. Daniel

**Affiliations:** Department of Cell Biology and Genetics, University of Lagos, Lagos, Nigeria; Division of Immunology, Institute of Infectious Disease and Molecular Medicine and Department of Pathology, University of Cape town, South Africa; Immunology Unit, Medical Laboratory Science Department, Nnamdi Azikiwe University, Nnewi Campus, Nigeria; Department of Veterinary Pathology, University of Ibadan, Ibadan, Nigeria; Public Health Biotechnology Unit, Institute of Child Health, University College Hospital, University of Ibadan, Nigeria

**Keywords:** Covid-19, SARS-CoV-2, Epitope Vaccine, Reverse Vaccinology, Molecular dynamics simulation

## Abstract

Developing an efficacious vaccine to SARS-CoV-2 infection is critical to stem COVID-19 fatalities and providing the global community with immune protection. We have used a bioinformatic approach to aid in the design of an epitope peptide-based vaccine against the spike protein of the virus. Five antigenic B cell epitopes with viable antigenicity and a total of 27 discontinuous B cell epitopes were mapped out structurally in the spike protein for antibody recognition. We identified eight CD8^+^ T cell 9-mers along with 12 CD4^+^ T cell 14-15-mer as promising candidate epitopes putatively restricted by a large number of MHC-I and II alleles respectively. We used this information to construct an *in silico* chimeric peptide vaccine whose translational rate was highly expressed when cloned in pET28a (+) vector. The vaccine construct was predicted to elicit high antigenicity and cell-mediated immunity when given as a homologous prime-boost, with triggering of toll-like receptor 5 by the adjuvant linker. The vaccine was characterized by an increase in IgM and IgG and an array of Th1 and Th2 cytokines. Upon *in silico* challenge with SARS-CoV-2, there was a decrease in antigen levels using our immune simulations. We therefore propose that potential vaccine designs consider this approach.

## Introduction

An unprecedented pneumonia disease outbreak was reported in late December 2019, after several deaths were recorded in Wuhan, China **[1]**. There was a rapid spread of the disease from the city of Wuhan to many countries including the United States with thousands infected and many dying within months of initial spread [**1, 2**]. On the 31^st^ of December 2019, the disease outbreak was traced to a novel strain of coronavirus **[1]**, which was later termed SARS-CoV-2 and the disease COVID-19 by the WHO **[3, 4]**. Reports showed that as of 16^th^ June 2020, there have been at least 440,421 confirmed deaths and more than 8,144,359 confirmed cases **[5]**. SARS-CoV-2 been identified as a new strain from group 2B Coronaviruses, with approximately 70% genetic similarity to SARS-CoV, from the 2003 outbreak **[4]**. The virus has a 96% similarity to a bat coronavirus, so it is widely suspected to originate from bats **[6, 7]**. The pandemic has resulted in travel restrictions and nationwide lockdowns in several countries and resulted in economic mayhem **[1]**. In the search for solutions, genome sequences have been deposited in several online repositories, and developing a universally available vaccine is critical.

Coronaviruses are group of viruses that generally causes disease in mammals and Aves. In the case of humans, coronaviruses are responsible for respiratory tract infections that can range from mild, such as cases of the common cold and low fever, and others that can be lethal, such as SARS, MERS, and COVID-19 **[4]**. Cases have shown that symptoms can vary depending on the host, where for example, upper respiratory tract disease has been found in chickens and diarrhea in cows and pigs [**6**]. The Coronavirus is grouped under the subfamily *Orthocoronavirinae*, in the family Coronaviridae, order Nidovirales **[5, 6]**. They are positive-sense single stranded RNA virus with nucleocapsid of helical symmetry. They are also enveloped with spike proteins and the genome size of many of the coronaviruses range from 27-34 kilobases, the largest among known retroviruses **[8]**.

The current zoonotic jump to humans is of concern as the upper respiratory tract infection can lead to fatal disease in some individuals. There is yet to be an effective vaccine to prevent or treat any human coronavirus infections.

Structural analysis has revealed that the Spike protein S1 attaches the virion to the cell membrane by interacting with ACE2 and CLEC4M/DC-SIGNR receptors. The Internalization of the virus into the endosomes of the host cell induces conformational changes in the S glycoprotein **[9]**. Proteolysis by cathepsin may unmask the fusion peptide of S2 and activate membranes fusion within host endosomes. Spike protein S2 mediates fusion of the virion and cellular membranes by acting as a class I viral fusion protein. Under the current model, the protein has at least three conformational states: pre-fusion native state, pre-hairpin intermediate state, and post-fusion hairpin state **[9]**. During viral and target cell membrane fusion, the coiled regions (heptad repeats) assume a trimer-of-hairpin structures, positioning the fusion peptide in close proximity to the C-terminal region of the ectodomain **[10]**. The formation of this structure appears to drive apposition and subsequent fusion of viral and target cell membranes. Spike protein S2’ acts as a viral fusion peptide which is unmasked following S2 cleavage occurring upon virus endocytosis **[10]**. The cellular receptor of SARS-CoV-2 and the receptor-binding domain (RBD) on the S protein has been identified **[11, 12]**. Previous studies have shown that the RBD in the S1 region plays a critical role in neutralizing antibody induction, angiotensin-converting enzyme 2 (ACE2) binding and virus entry **[13]**. Depletion of RBD-specific antibodies from sera significantly reduced serum-neutralizing capability, indicating that this domain is dominant in neutralizing antibody induction **[14]**. The pivotal role of the S protein in viral infection has made it a top candidate for vaccine production. There are currently over 90 SARS-CoV-2 vaccine candidates **[1]** and an epitope-based vaccine may provide a useful complimentary approach that would steer immunity to immunogenic epitopes on the S protein. With the immuno-informatics, it is now possible to evaluate the immunogenic properties of proteins via computational methods (*in silico*) with high efficiency and confidence [**15–17**]. We used such an approach to design an epitope peptide-based vaccine against SARS-CoV-2 spike glycoprotein and then *in silico* mimic the range of responses in a prime-boost scenario.

## Materials and Methods

### Data retrieval, Structural and Physiochemical Analysis of SARS-CoV-2 Spike Protein

The protein sequence from different geographical regions was retrieved from the NCBI repository with their corresponding accession numbers: Wuhan, China (Genbank ID: QHD43416.1), Japan (Genbank ID: BCA87361.1), California, USA (Genbank ID: QHQ71963.1), Washington, USA (Genbank ID: QHO60594.1), and Valencia, Spain (Genbank ID: QIQ08790.1). The protein structure of the SARS-CoV-2 spike (PDB: 6VSB) was downloaded from the protein data bank. The physiochemical properties of the protein sequence such as the GRAVY (Grand average of hydropathicity), half-life, molecular weight, instability index, aliphatic index, and amino acid atomic composition was bio-computed via an online tool Protparam (http://web.expasy.org/protparam/) **[18]**.

### Prediction of B Cell Linear and Discontinuous Epitopes

The Bepipred server from the Immune-Epitope-Database and Analysis-Resource (IEDB) database was used for this prediction was used to identify B cell linear epitopes [**19**]. Bepipred-2.0 is based on a random forest algorithm trained on epitopes annotated from antibody-antigen protein structures [**19**]. This method is superior to other available tools for sequence-based epitope prediction with regards to both epitope data derived from solved 3D structures and a large collection of linear epitopes downloaded from the IEDB database [**19**]. The following criteria such as the specificity at 75% and 14-15 mers (residues) was assumed to bind to MHC. Several conditions such as antigenicity, accessibility of surface, flexibility, hydrophilicity is imperative for the prediction of the B cell linear epitope. These conditions are taken into consideration when making predictions with the Bepipred linear epitope prediction and Parker hydrophilicity prediction algorithms.

SVMTriP (http://sysbio.unl.edu/SVMTriP/) was also used in the prediction of the B cell linear epitopes. The SVMTriP is a Support Vector Machine method used to predict linear antigenic epitopes which combine the Tri-peptide similarity and Propensity scores (SVMTriP). Application of SVMTriP to non-redundant linear B-cell epitopes extracted from IEDB achieved a sensitivity of 80.1% and a precision of 55.2% with five-fold cross-validation. For antigenicity testing, these epitopes were subjected Vaxijen 2.0 **[20]**. We further predicted the discontinuous epitopes which possesses greater attributes than the linear epitopes and discovery of discontinuous B-cell epitopes is a major challenge in vaccine design. Previous epitope prediction methods have mostly been based on protein sequences and are not very effective. Therefore, the DiscoTope server was used to predict the surface accessibility and amino acids that form discontinuous B cell epitopes found from X-ray crystallography of antigen/antibody protein buildings. Pymol was utilized to examine the positions of forecast epitopes on the 3D structure of SARS-CoV-2 protein [**21**].

### Prediction of epitopes restricted by class I Human Leukocyte Antigen (HLA) CD8+ (CTL) and class II HLA CD4+ T cells (HTL)

For *de novo* prediction of Covid-19 spike glycoprotein CD8^+^ T cell epitopes (peptides), we used IEDB MHC I binding prediction algorithms (http://tools.iedb.org/mhci). This method integrates the prediction of epitopes restricted to a large number of MHC class I alleles and proteasomal C-terminal cleavage, using artificial neural network application. For better predictive accuracy, other software such as artificial neural network (ANN), stabilized matrix method (SMM), MHC binding energy covariance matrix (SMMPMBEC), NetMHCpan, pickpocket, and NetMHCstapan, were adopted for this purpose. All of these predictive tools are archived on the IEDB (Immune-Epitope-Database and Analysis-Resource) database with mathematical threshold before best-fit epitopes are selected from each online server. To predict the CD4^+^ T cell epitopes (peptides), we used the MHC II binding predictions tool (http://tools.iedb.org/mhcii/) found in the IEDB database. First, we selected the epitopes whose binding diversities with the different HLA serotypes were higher, and we further subjected these epitopes to Vaxijen 2.0 server to test for their antigenicity at a recommended threshold of 0.7. We also considered for further analysis, by subjecting the top-scoring predicted epitopes from each tool that had been predicted by five or more different methods and submitted them to IEDB T cell Class I Immunogenicity predictor (http://tools.iedb.org/immunogenicity/). Results were given in descending score values. However, the table can also be sorted by clicking on individual column headers. The higher score indicates a greater probability of eliciting an immune response.

### Profiling of the selected T cells Epitopes

Following the selection of HLA-restricted CD8^+^ and CD4^+^ T cell epitopes, critical features such as peptide toxicity predicted from the ToxinPred server (http://crdd.osdd.net/raghava/toxinpred/), allergenicity, predicted from AllergenFP 1.0 and digestion predicted from protein digest server were made. All of these criteria were considered before the final selection of the T cell epitopes. Epitopes with no toxicity were selected. Antigenicity testing was conducted through the Vaxijen v2.0 server (http://www.ddgpharmfac.net/vaxijen/VaxiJen/VaxiJen.html) [**20**], which operate based on auto- and cross-covariance transformation of the input protein sequence into uniform vectors of principal amino acid properties. The antigenicity index is generated at a threshold of 0.7.

### Epitope Conservancy in related SARS-CoV-2 Spike protein from different geographical locations

Conservation analysis of selected epitopes is the fraction of a protein sequence that contains the epitope, while the identity is the degree of correspondence (similarity) between the sequences. The degree of epitope conservancy was computed within the SARS-CoV-2 spike glycoprotein sequence and set at a given identity level of 100 using the IEDB conservation-analysis-tool.

### The HLA-A 02*01 Allelic Affinity of the CD8^+^ T cell epitopes

The Molecular docking of the antigenic epitopic peptides was conducted with the alleles they were mostly restricted to, of which the HLA-A 02*01 allele was included. The X ray crystal structure of the molecule was retrieved from the protein data bank (PDB: 4U6Y) and dock with the epitopes. The refined binding free and dissociation energies were determined from the docked complex.

### Population Coverage Analysis of CD8+ and CD4+ T cell epitopes

The selected epitopes from the HLA class I and class II families together with their respective binding leukocyte antigens were subjected to IEDB Population Coverage tool (http://tools.iedb.org/population/). This calculated the distribution or fraction of individuals predicted to show a response to the selected epitopes with known HLA background. The tool also computes the average number of epitope hits/HLA allele combinations recognized by the entire population and the maximum and minimum number of epitope hits recognized by 90% of the selected population. The HLA genotypic frequencies are calculated and T cell epitopes queried based on the area, ethnicity and country. The entire world population was selected, followed by subcontinents and countries. For countries like Nigeria and Ghana with no deposited information on the IEDB database, they were included as part of the West African population.

### Designing of Multi-Epitope Vaccine Construct

Selected antigenic epitopes were scrutinized to determine which could potentially induce different Th1 and Th2 cytokines. Those with this attribute were selected for the vaccine construct. To construct a multi-epitope vaccine, we finally selected CTL, HTL, and B cell linear epitopes that were linked together with the help of AAY, GPGPG, and KK linkers, respectively. To boost the immunogenic profile of the selected profile epitopes, an adjuvant would be required. The outer membrane protein A (OmpA) (GenBank: AFS89615.1) was retrieved for this purpose, because its serves as agonist to the human immune receptor by interacting with antigen presenting cells **[22]**. The adjuvant was putatively added through the EAAAK linker, with the B and HTL epitopes which were linked together through the GPGPG linkers. These complexes were subsequently added to the CTL epitopes through the AAY linkers. The tag (6xHis-tag) was added at the C terminal end of the vaccine construct. The 6xHis-tag is one of the simplest and most widely used purification tags, with six or more consecutive histidine residues. These residues readily coordinate with transition metal ions such as Ni^2+^ or Co^2+^ immobilized on beads or resin for purification **[23]**. The intrinsic solubility properties of the vaccine peptide were conducted using the CamSol tool, which yields a solubility profile where regions (residues) with scores greater than 1 signifies soluble regions, while scores lesser than −1 represents poorly soluble regions **[24]**. An overall score is generated for the entire sequence, as these amino residue scores are ranked based on their level of solubility.

### Structural Modelling, Assessment, and Validation

All the predicted peptides 3D structures were modelled via PEPFOLD server at RPBS MOBYL portal [**25**]. PEP-FOLD is a *de novo* approach aimed at predicting peptide structures from amino acid sequences [**25, 26, 27**]. This method, based on structural alphabet SA letters to describe the conformations of four consecutive residues, couples the predicted series of SA letters to a greedy algorithm and a coarse-grained force field [**25, 26**]. The predicted models are in cluster ranks which are defined according to their scores. The cluster representatives correspond to the models of the clusters having the best scores, i.e. with the lowest sOPEP energy (representing the highest tm value) **[25]**. The PSIPRED v4.0 server was adopted for the prediction of the vaccine’s secondary structure **[28]**, while the Swiss dock online tool was used for the tertiary structure prediction of both the vaccine construct and the human HLA class II histocompatibility antigen, DR alpha chain. To validate the generated protein structure, Procheck online tool together with Ramachandran plot analysis was generated **[29]**. The plot analysis was able to show the allowed and disallowed dihedral angles psi (ψ) and phi (□) of an amino acid which is calculated based on van der Waal radius of the side chain. The corresponding percentage value of both the allowed and disallowed region of the separate plots of glycine and proline residues of the modeled structure was generated.

### Molecular Docking Studies

One of the best ways to access the immune response mounted by the epitopes is by studying their binding affinity characterizing their molecular interaction with the human HLA class I and I molecules. The binding pockets on the HLA-class I and II molecules and the human immune receptor (TRL5) was predicted using the CASTp server (http://sts.bioe.uic.edu/castp/). The CASTp server provides comprehensive and detailed quantitative characterization of topographic features of a protein. The geometric modeling principle involves the calculation strategy of alpha-shape and discrete-flow methods that are applied to the protein binding site, also the measurement of pocket-size by the program **[30]**. The protein pocket atom is identified and then the volume and area are calculated **[31, 32]**. The program also identifies the atoms forming the rims of pocket mouth, computes how many mouth openings for each pocket, predict the area and circumference of mouth openings, finally locates cavities and calculates their size. The secondary structures were calculated by DSSP **[30, 32]**. The predicted structure of the HLA class I and class II allele protein were utilized for molecular docking analysis with the selected epitopes (peptides) to evaluate their binding affinities. The protein structure was chemically manipulated by expulsion of water and ligand molecules. For the peptide-protein interaction, HPEPDOCK (http://huanglab.phys.hust.edu.cn/hpepdock/) was utilized for this purpose [**33**]. It uses a hierarchical algorithm and instead of running lengthy simulations to refine peptide conformations, HPEPDOCK also considers peptide flexibility. UCSF Chimera and Pymol tools were utilized to produce figures of docked complexes. ZDOCK server **[34]** was adopted for the molecular docking between the multiple epitope vaccine peptides and the human immune receptor (PDB: 3J0A). ZDOCK is based on the rigid-body docking program that predicts protein-protein complexes and symmetric multimers. ZDOCK achieves high predictive accuracy on protein-protein docking benchmarks, with >70% success in the top 1000 predictions for rigid-body **[35]**.

### Molecular Dynamics Simulation Studies

The biological molecules in a solution of the peptide vaccine construct was studied, using the small -and wide-angle X-ray scattering (SWAXS) **[36]**. The generated curves require accurate prediction from the structural model. The predictions are complicated by scattering contributions from the hydration layer and by effects from thermal fluctuations. The MD simulations provide a realistic model for both the hydration layer and the excluded solvent, thereby avoiding any solvent-related fitting parameters, while naturally accounting for thermal fluctuations **[36]**. To determine the protein compactness, the radius of gyration of the biomolecule through the Guinier analysis was also conducted. The interacting complex between the vaccine and the toll-like receptor (PDB: 3J0A), was thoroughly accessed based on the existing coordinates between the docked protein complex. Parameters considered were the deformability, B factor, eigenvalues associated with the normal mode which represents the motion stiffness. Its value is directly related to the energy required to deform the structure. The lower the eigenvalue, the easier the deformation. The covariance matrix was also considered for the simulation. It indicates the coupling between the pairs of residues. The correlation matrix is computed using the Cα Cartesian coordinates. The elastic network of the docked complex was also computed **[36]**.

### *In Silico* Codon Adaptation and Cloning

For the maximum expression of the vaccine in the host, a codon optimization was conducted. This was done using the Java Codon Adaptation Tool (JCat), with the aim of boosting the vaccine translational rate in *E. coli* K12. Restriction enzymes cleavage sites, prokaryote ribosomal binding site, and finally rho-independent transcription termination, were all avoided during the option selection. Codon adaptation index (CAI) value and GC content of the adapted sequence was obtained and compared with the ideal range. The obtained refined nucleotide was cloned into the pET28a (+) vector, utilizing the SnapGene 4.2 tool.

### Immune Simulation of the Chimeric Peptide Vaccine

The entire predicted conjugate vaccine peptide was accessed for their immunogenicity and immune response attributes using the C-ImmSim online server (http://150.146.2.1/C-IMMSIM/index.php) **[37]**. The server uses a machine-learning basis in predicting the epitopes and the associated immune interactions. It automatically simulates three anatomical compartments which include: (i) bone, where the hematopoietic stem cells are stimulated and myeloid cells are produced, (ii) the lymphatic organ and (iii) the thymus where naive T cells are selected to avoid autoimmunity. Three injections containing the designed peptide vaccine was administered at an interval of four weeks. From the default parameters, each time step were positioned at 1, 84, and 168 meaning that each time step is 8hours and time step 1 is the injection administered at time zero. So, three injections were administered at four weeks interval. However, eight injections were administered four weeks apart to stimulate repeated exposure to the antigen. In this scenario, the T cell memory will undergo continuous assessment. The Simpson index was graphically interpreted from the plot analysis **[37]**.

## Results

### Linear and Discontinuous B cell Epitopes

There were 5 promising linear B cell epitopes with non-allergenic attributes. The peptide “VRQIAPGQTGKIAD” comparatively had the highest antigenic index than the other predicted B cell epitope candidates. A characteristic non-toxic peptide attribute makes the selected antigenic epitopes safe for vaccine design. The antigen conservancy of the epitopes across the retrieved spike protein from different geographical locations was 100% [**Table 1**].

**Table 1:** B cells linear epitopes of SARS-CoV-2 Spike glycoprotein and their immunogenic properties.

| No. | Start | End | Peptide | Length | Antigenicity | Toxicity | Allergenicity | Conservancy |
| --- | --- | --- | --- | --- | --- | --- | --- | --- |
| 1 | 407 | 420 | VRQIAPGQTGKIAD | 14 | 1.261 | Non-toxic | Non-allergen | 100 |
| 2 | 18 | 32 | LTTRTQLPPAYTNSF | 15 | 0.79 | Non-toxic | Non-allergen | 100 |
| 3 | 4 | 18 | FLVLLPLVSSQCVNL | 15 | 0.8302 | Non-toxic | Non-allergen | 100 |
| 4 | 749 | 762 | SNLLLQYGSFCTQL | 14 | 0.800 | Non-toxic | Non-allergen | 100 |
| 5 | 1056 | 1069 | PHGVVFLHVTYVPA | 14 | 0.806 | Non-toxic | Non-allergen | 100 |

**NB:** All of the selected B cell are tagged B1-B5.

Graphically, using the Kolaskar and Tongaonkar antigenicity scale, Emini surface accessibility, and the Chou and Fasman beta-turn predictions, regions with viable immunogenic properties were determined. The scale was able to show the favorable regions across the protein that are potentially antigenic [**supplementary data: Figure s1a-s1c**]. The resulting B cell linear epitopes were mapped out from the spike protein **[Figure 1a]**. The predicted discontinuous epitopes were selected from the entire protein chain component (A, B, and C) of the virus spike protein (PDB: 6VSB), and ranked based on their propensity scores. A total of 27 discontinuous epitopes were mapped out from the protein structure as shown in **Figure 1b**. Across the discontinuous epitopes of the protein chain components, the maximum contact number was 10, and the least was 7. The chain C component of the spike protein had a higher number of contact residues [**supplementary data: S-Table 2]**.

**Figure 1a:**
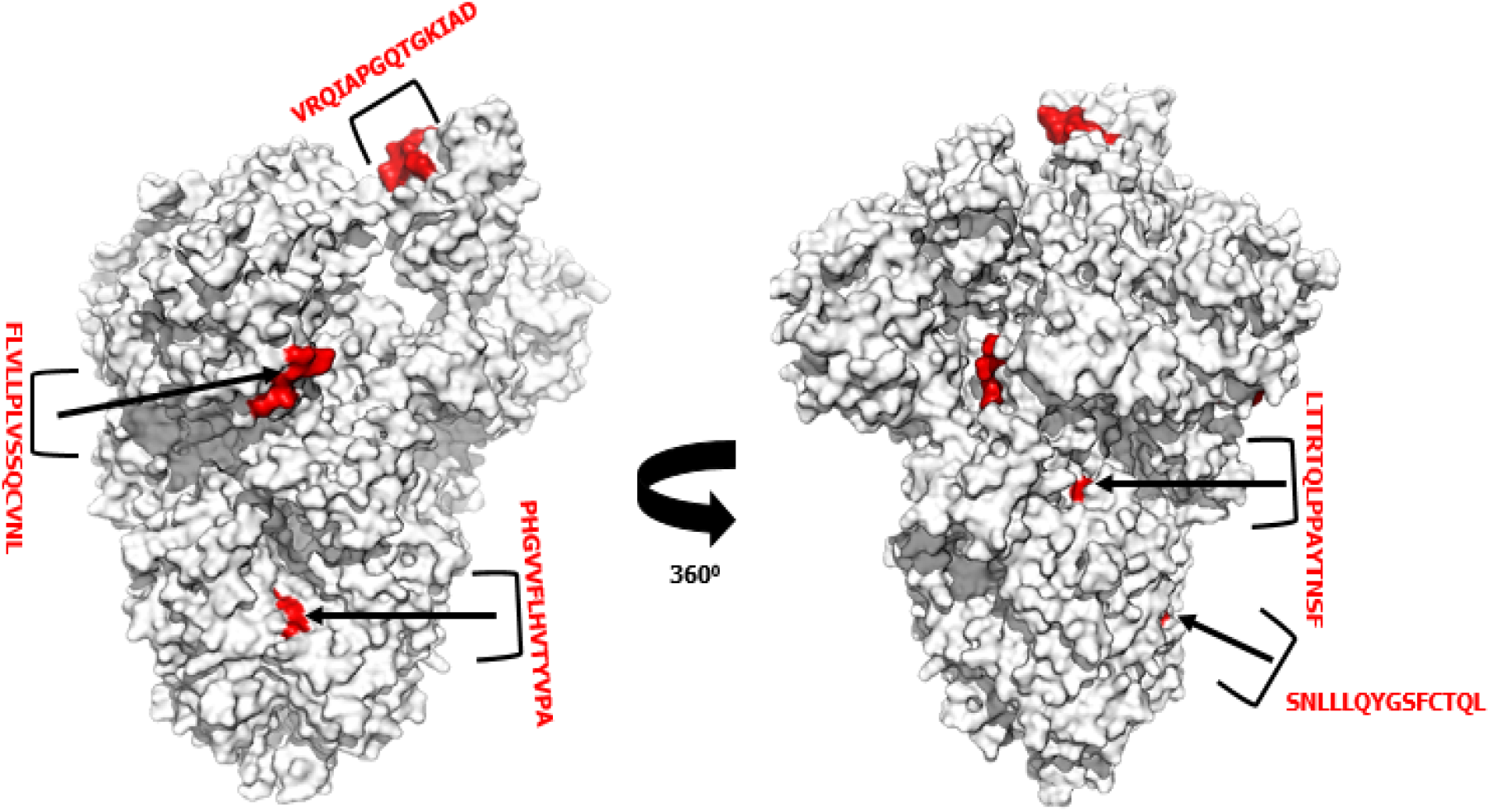
SARS-CoV-2 crystal spike glycoprotein showing the mapped-out B cell linear epitopes. The linear epitopes are highlighted in red.

**Figure 1b:**
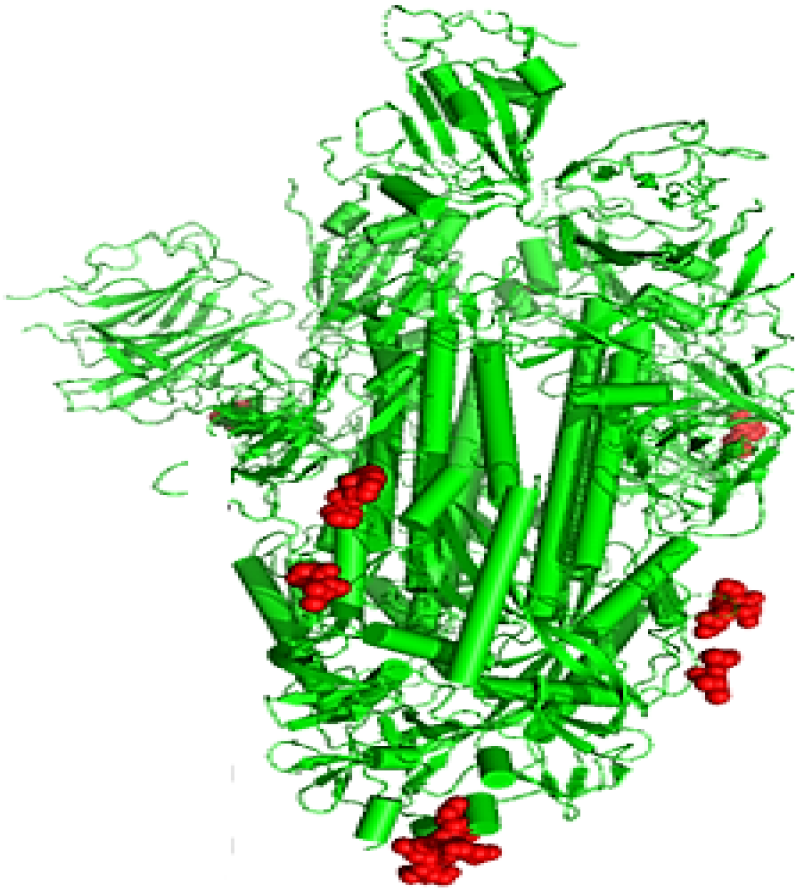
Site of the predicted discontinuous B cell epitopes overlaid on the crystal structure of SARS-CoV-2 envelope S protein. The discontinuous epitopes are shown in clusters of red spheres.

### The CTL and HTL Epitopes

The spike protein sequence was scanned across multiple HLA class 1 alleles. The peptides were selected based on their percentage rankings and the number of alleles they potentially bind to. Additionally, the peptides were subjected to antigenicity test using Vaxijen 2.0. Based on the antigenicity scores, 16 epitopes were selected for the next stage of screening. The most important peptides are those with the capacity of binding with a higher number of HLA class I molecules and showing a non-allergenic attribute. Before vaccine design can be considered, the allergenicity prediction is crucial, as there is a possibility of vaccine candidates eliciting a Type II hypersensitivity reaction. Allergen 1.0 online was adopted for this analysis and the allergenicity scores show that these epitopes were non-allergenic. The non-toxicity attribute of the peptides also makes them suitable for vaccine production. Eight peptides were allergenic and eight were also non-allergenic.

The non-allergenic peptides were: GAEHVNNSY which is putatively restricted to HLA-A*01:01, HLA-B*15:01, and HLA-C*02:02, KTSVDCTMY attaches to 5 alleles: HLA-A*01:01, HLA-B*15:01, HLA-C*02:02, HLA-A*03:01, HLA-A*30:01, TTEILPVSM would be able to bind to 3 alleles: HLA-A*01:01, HLA-C*02:02, HLA-C*01:02. Other selected epitopes had similar putative restricted attachments such as “ILDITPCSF” with an antigenicity score of 1.184 and potentially attaches to 7 alleles: HLA-A*01:01, HLA-C*04:01, HLA-C*02:02, HLA-C*01:02, HLA-B*15:01, HLA-A*02:01, HLA-B*13:01. The peptide GVYFASTEK would be able to bind to 3 alleles: HLA-A*03:01, HLA-A*30:01, HLA-C*02:02, GVYYHKNNK, ASANLAATK, VLKGVKLHY had the same similar attribute of binding to two, three and five alleles respectively [**Table 2**].

**Table 2:**
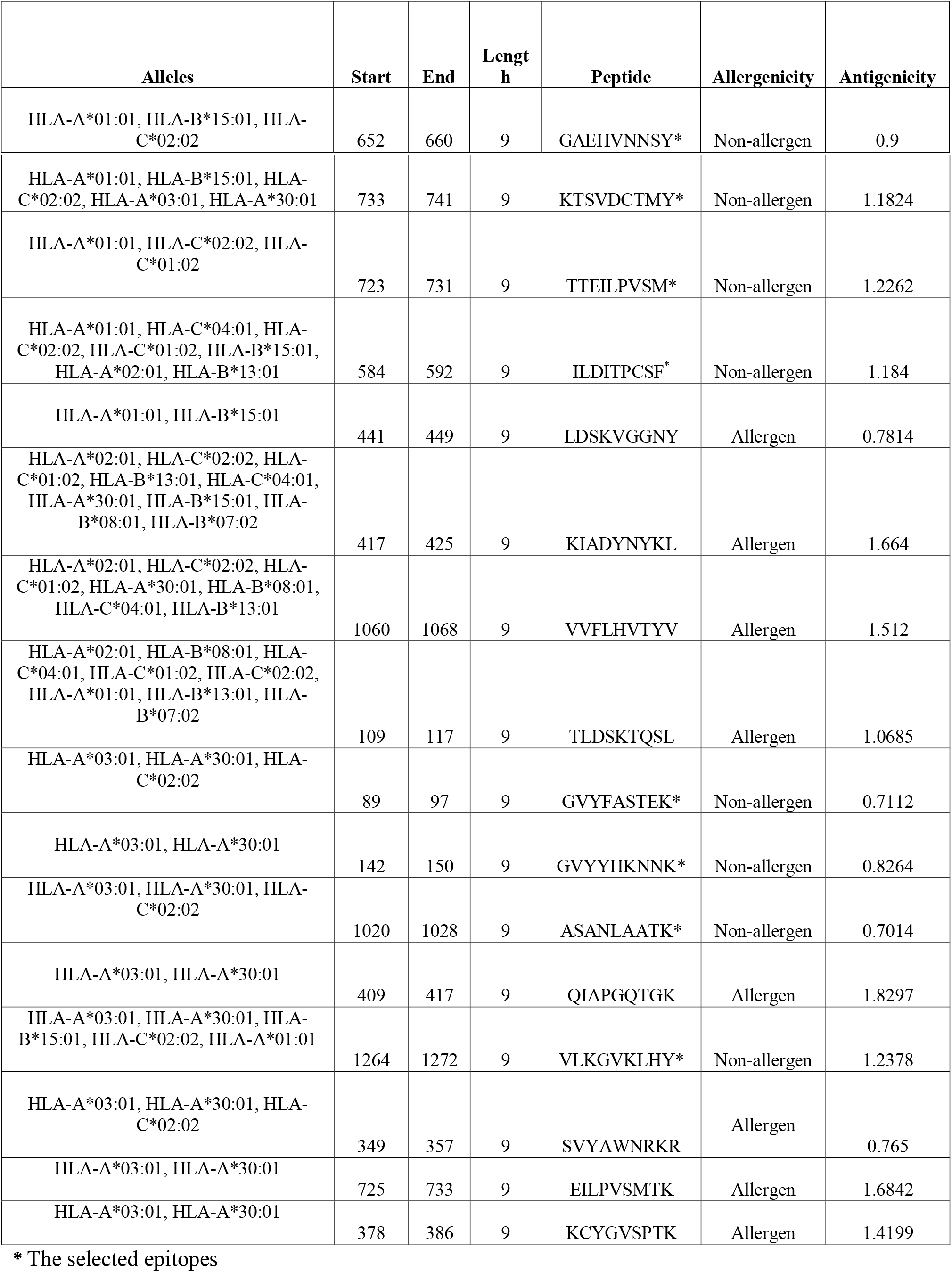
MHC-I T cell epitopes of SARS-CoV-2 spike glycoprotein.

For the HLA class II T cells epitopes, the spike protein sequence was also scanned through a large number of the MHC-II alleles. Twelve epitopes were selected based on their antigenic properties. All of these selected non-allergenic epitopes are capable of eliciting an immune response by inducing either or all of IFN-γ, IL-4 and IL-10 cytokines. The peptide “GYFKIYSKHTPINLV” was the only candidate to induce all of the three cytokines, which was intriguing. Selection of these epitopes was also centered on their putative bindings to a large number of MHC-II alleles. The peptide “FAMQMAYRF”, with an antigenicity score of 1.0278, attaches to 8 HLA alleles: -DRB1*01:02, -DRB1*01:04, -DRB1*01:03, -DRB1*01:01, -DRB1*01:05, -DRB1*07:01, -DRB1*04:01, and -DRB1*03:01. The epitope “FRVQPTESI”, with an antigenicity score of 0.9396, is also restricted to 8 HLA alleles: -DRB1*04:01, - DRB1*01:01, -DRB1*01:05, -DRB1*07:01, -DRB1*01:03, -DRB1*01:02, -DRB1*03:01, and - DRB1*01:04. The HLA-DRB1 is the most common and versatile MHC-II molecule. The entire putative attachments of the selected epitopes are summarized **[Table 3]**. The conservancy level of the epitopes across the retrieved spike protein sequences from different geographical location was 100%.

**Table 3:**
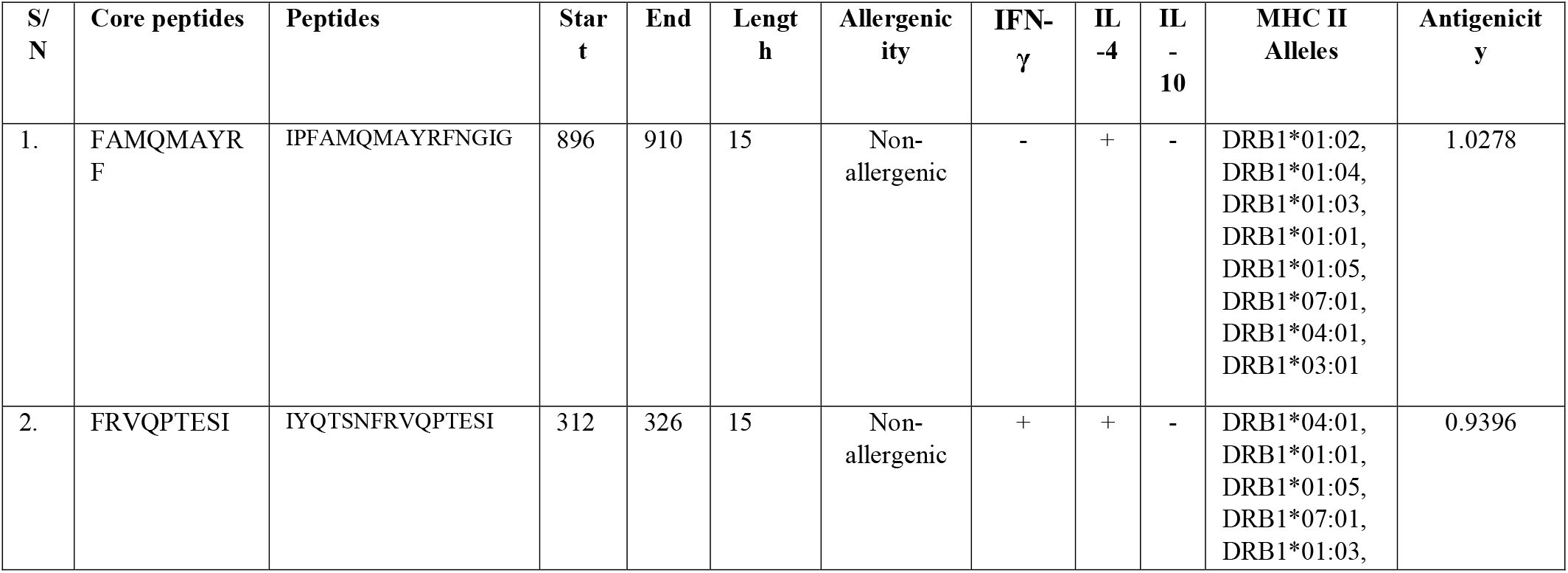
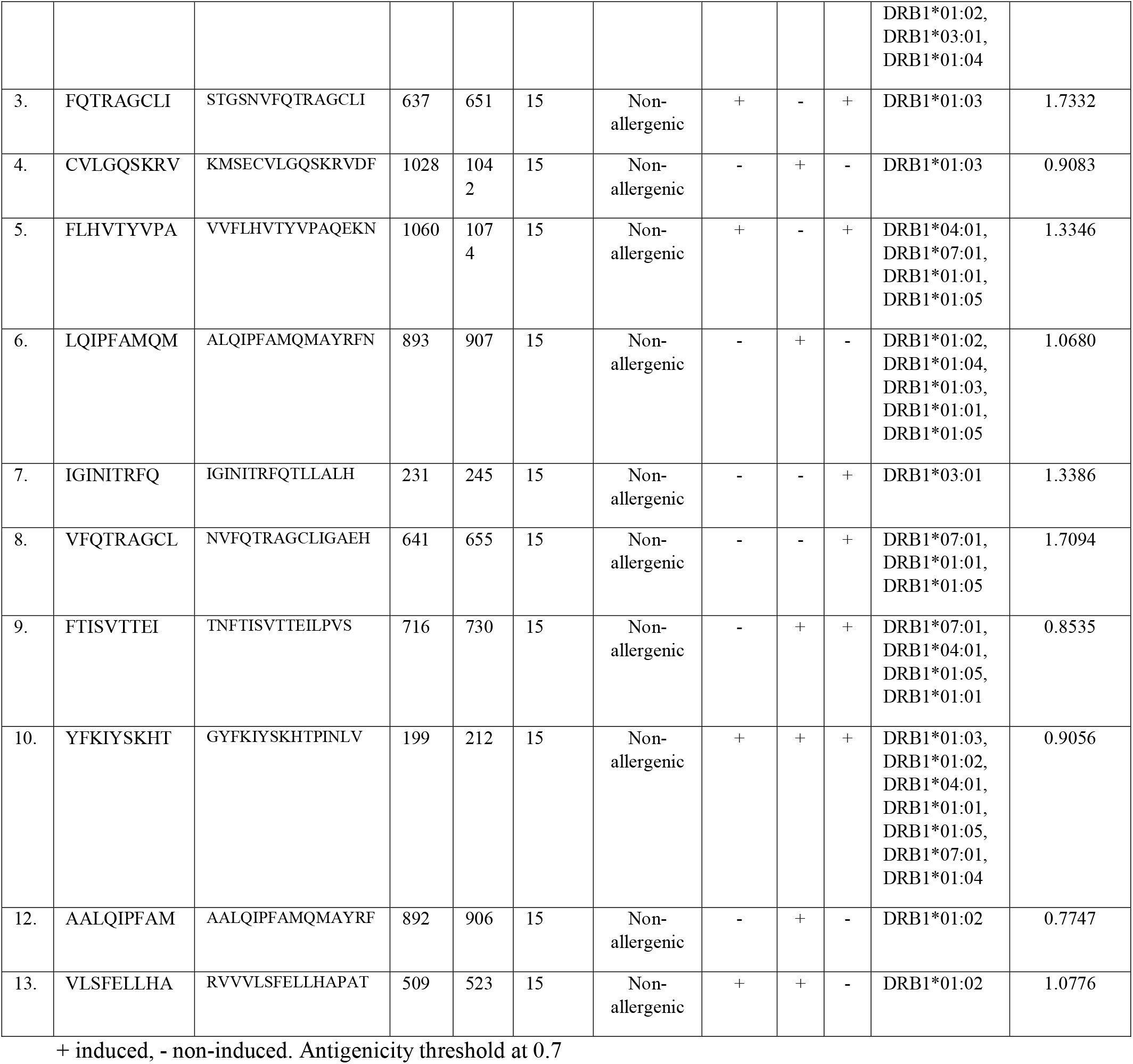
MHC class II T-cell Epitopes of SARS-CoV-2 spike glycoprotein

### Population Coverage of the CTL and HTL Epitopes

HLA allelic distribution differs among diverse geographical regions and ethnic groups around the globe. It is therefore imperative to consider the population coverage in designing a viable epitope-based vaccine relevant for global populations. The selected CD8^+^ T cell epitopes exhibited a higher individual percentage cover when queried with the entire world population. The HLA hits across the entire population revealed that approximately 81% of the world individuals are capable of responding to a median of 3 CTL epitopes [**Table 4**].

**Table 4:**
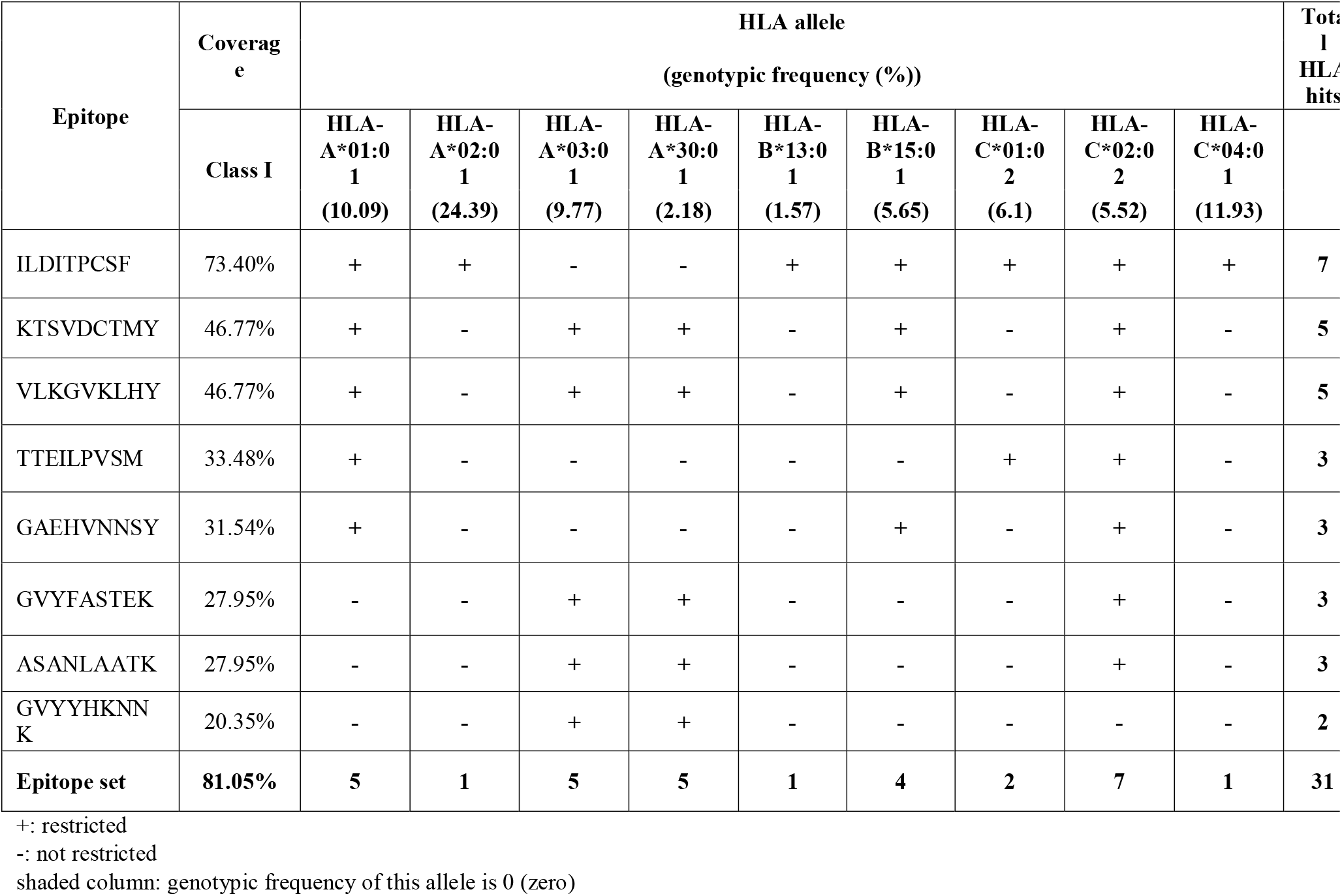
Coverage of individual epitope (MHC class I) in the world

However, the population coverage for the CD4^+^ T cell epitopes was comparatively lower compared to the CD8^+^ T cell epitopes, with average population coverage of 55.23% and recognition of a median of 2 epitopes [**Table 5**].

**Table 5:**
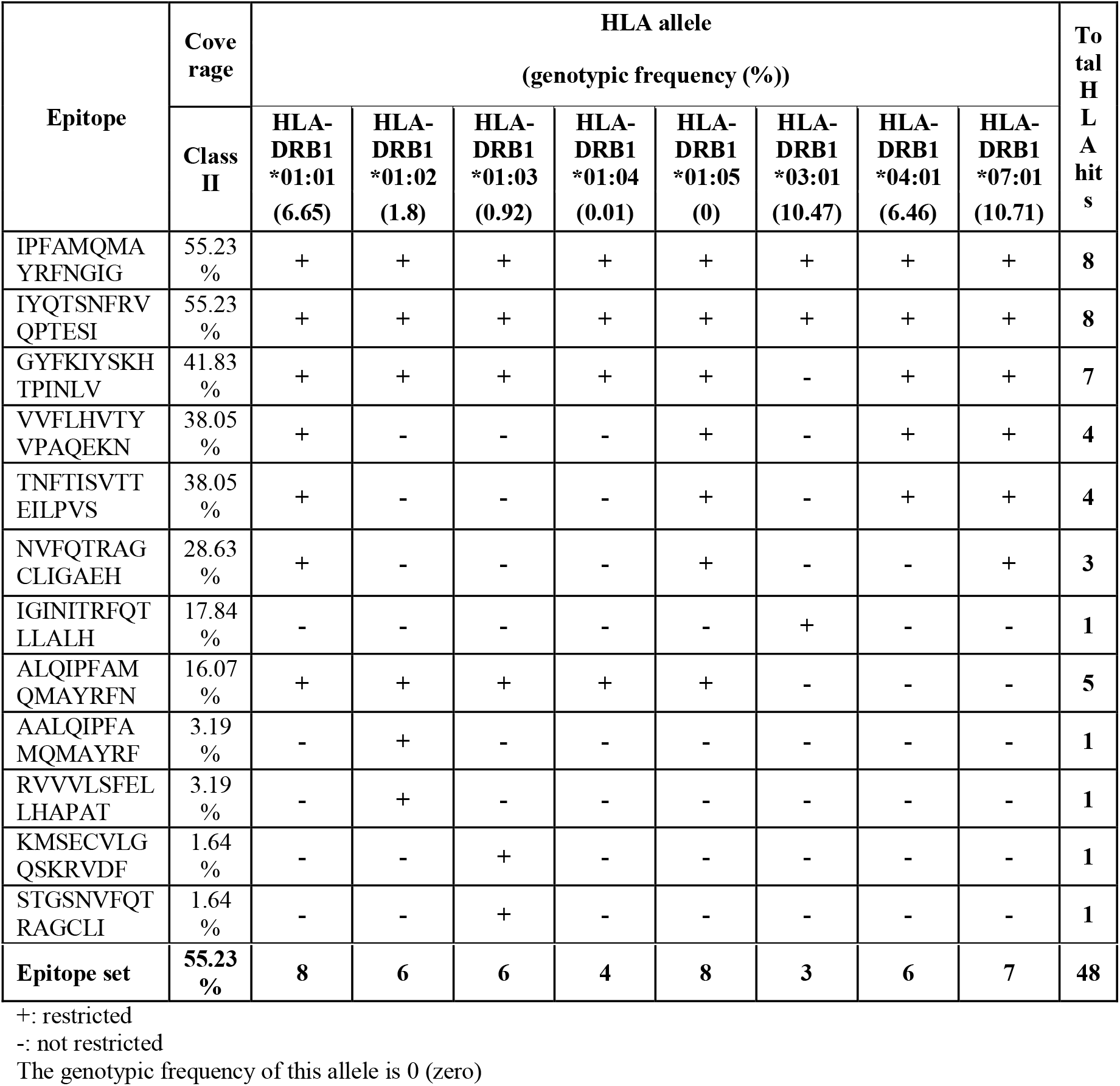
Coverage of individual epitope (MHC class II) in the world

Based on the selection of the continents and countries, the European populace would likely show a significant response to the selection of putative HLA class I restricted epitopes. England, France, United States, Italy, and Oceania had the highest population coverage of 92.31%, 85.75%, 82.22%, 80.39%, and 75.07% respectively, while the Pakistan population had the lowest population coverage at 35.8%. The population cover for the MHC class II epitopes in contrast to the MHC class I epitopes was considerably lower. The striking observation was 0% coverage exhibited by the Pakistan population **[Figure 2]**.

**Figure 2:**
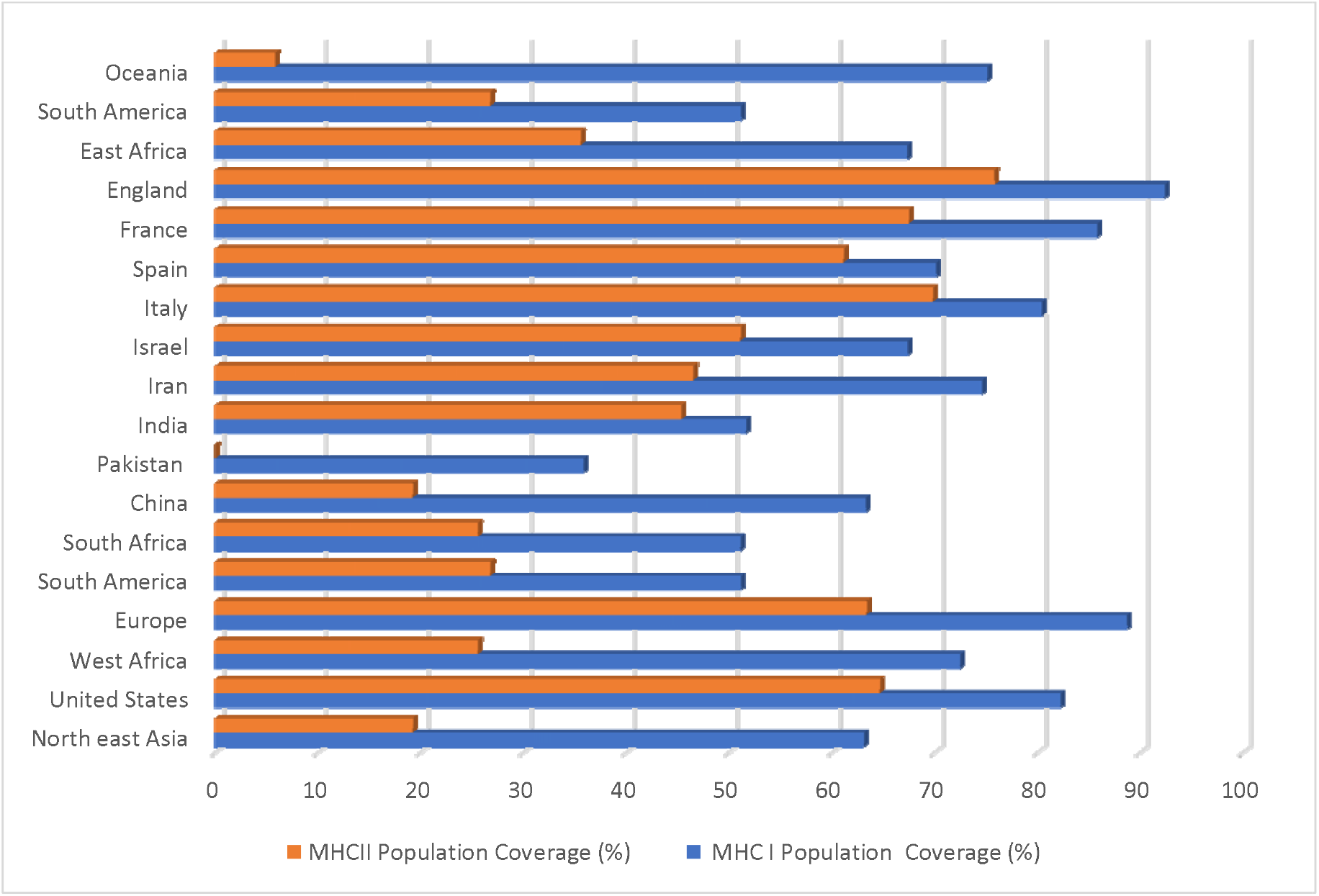
Selected continents and countries’ coverage of the MHC class I and II T cell epitopes.

### Binding Orientations of the CTL and HTL Epitopes in HLA-A*02:01 and HLA-DRB1*01:01 Groove

The selected CTL and HTL antigenic epitopes were docked individually with the alleles they were highly restricted to, which was HLA-A*02:01 for the CTL epitopes and HLA-DRB1*01 for the HTL epitopes. The differential binding patterns of the CTL epitopes were examined [**Figure 3ai-3aix**]. Major class histocompatibility class II amino acid sequences are highly polymorphic within a population, and correlate with individual differences in response to infectious agents and vaccines. It is therefore expedient to structurally examine how the CD4+ epitopes recognize peptide fragments of antigens that lie in the antigen groove of the MHC-II protein. The protein structure of the retrieved human HLA class II histocompatibility antigen, DRB1 beta chain (human leukocyte antigen DRB1, HLA-DRB1*01:01) (PDB: 1AQD) was retrieved for the molecular docking of the HTL epitopes because it was the most occurring allele that the peptides were restricted to. The HTL epitopes with good population cover were chosen for molecular docking with HLA-DRB1*01:01.

**Figure 3a-e:**
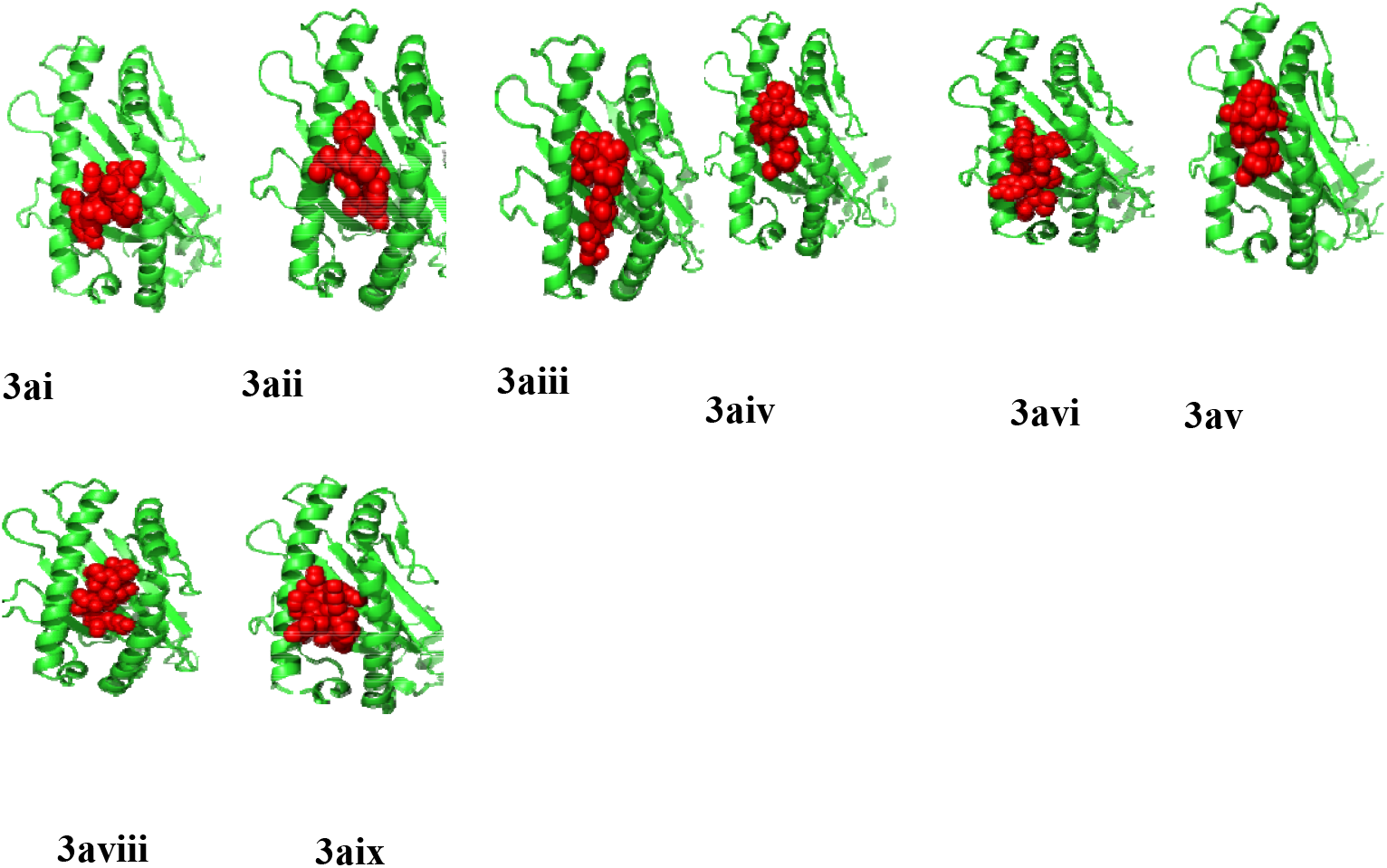
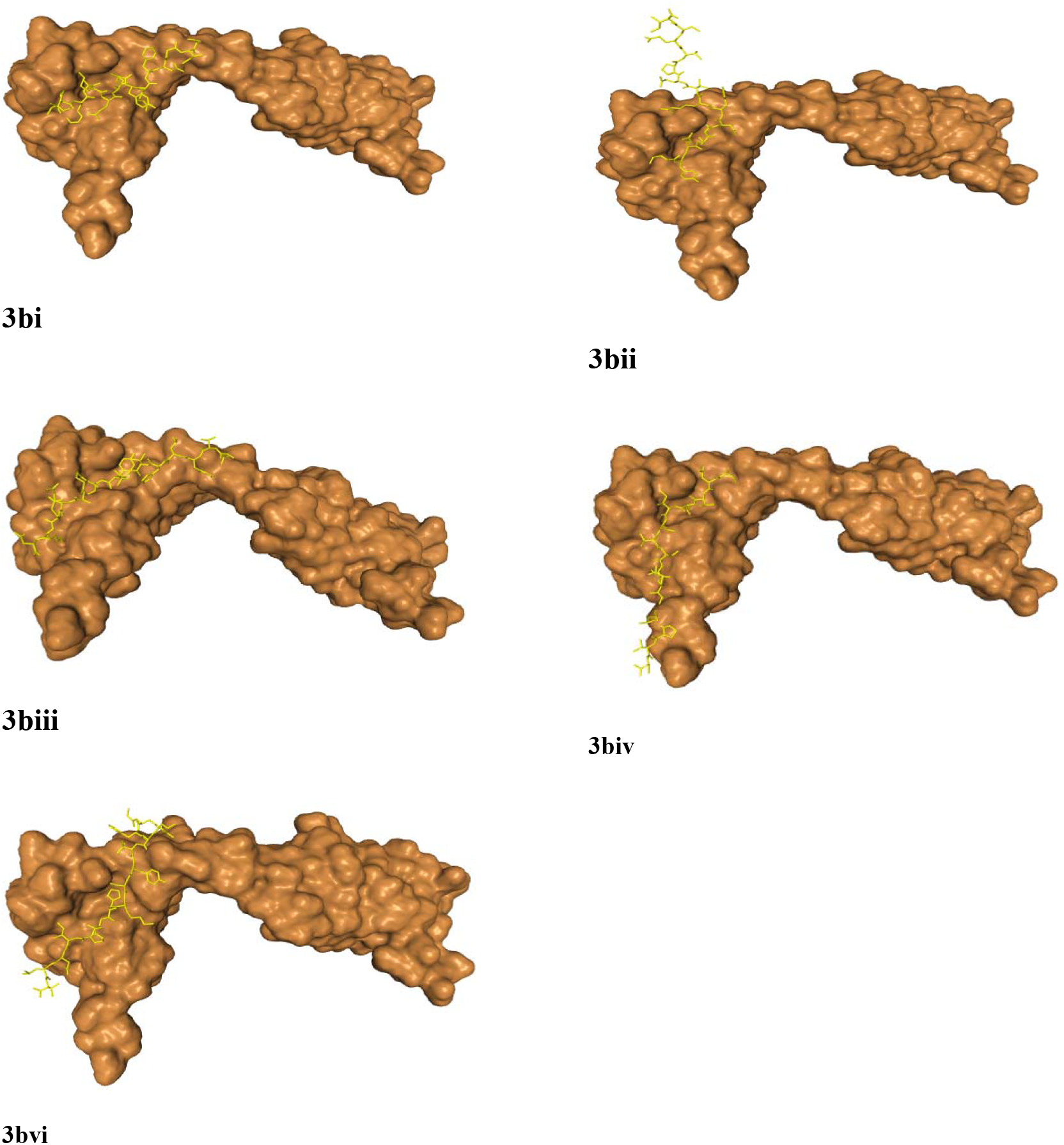
Differential binding of the HTL epitopes with HLA-DRB1*01:01. The MHC protein is displayed in surface brown and the epitopes are highlighted in yellow licorice.

The peptides: IPFAMQMAYRFNGIG, IYQTSNFRVQPTESI, VVFLHVTYVPAQEKN, TNTTISVTTEILPVS, and GYFKIYSKHTPINLV, were the selected epitopes. The binding free energy characterizing the HLA-DRB1 antigenic binding groove and the interacting epitopes alongside the corresponding amino residues was evaluated. The epitopes exhibited different binding pattern with the MHC class II groove. Few of the binding peptides had a flanking region outside the groove. Amino acids that are outside of the “core” peptide region extends out of the open MHC-II binding groove forming the peptide flanking regions at both the N- and C-terminus **[Figure 3bi-3biv]**. The epitope “IYQTSNFRVQPTESI” had the most extensive flanking non-binding region with some of part of the peptide protruding completely out of the groove. IPFAMQMAYRFNGIG had the best binding free energy score, with TNTTISVTTEILPVS with the least binding energy **[Table 6]**.

**Table 6:**
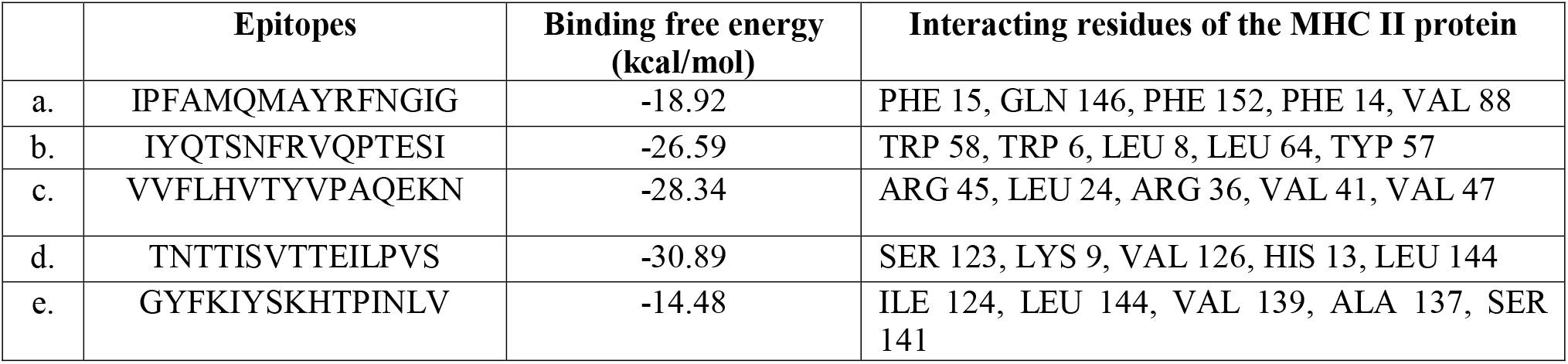
Docking properties of HLA-DRB1*01:01 restricted epitopes

### Construction of the Peptide Vaccine

The multiple epitope peptide vaccine consists of 553 amino residues from 25 selected antigenic B and T cells epitopes, covalently linked with an immuno-adjuvant [**Figure 4a]**. The tertiary structure of the multiple epitope vaccine was also obtained **[Figure 4b]**, and the structural validation was assessed using ProSA-web which predicts the overall quality of the model indicated in the form of z-score. If the z-scores of the predicted model are outside the range of the characteristic for native proteins, it indicates the erroneous structure. The Z-score was −2.32 for the vaccine predicted model indicating a relatively good model [**Figure 4c**]. Before the addition of the OmpA protein adjuvant, the conjugated vaccine was highly antigenic with a score of 0.8 after subjecting it to Vaxijen server, signifying that the vaccine is viable at inducing cellular and humoral immune response without the aid of an adjuvant. However, an adjuvant was added to further boost the immunogenic properties to 0.85. A structural appraisal of the secondary structure of the vaccine revealed 14% alpha helix, 41% beta strand and the disordered region was 17%.

**Figure 4a.**
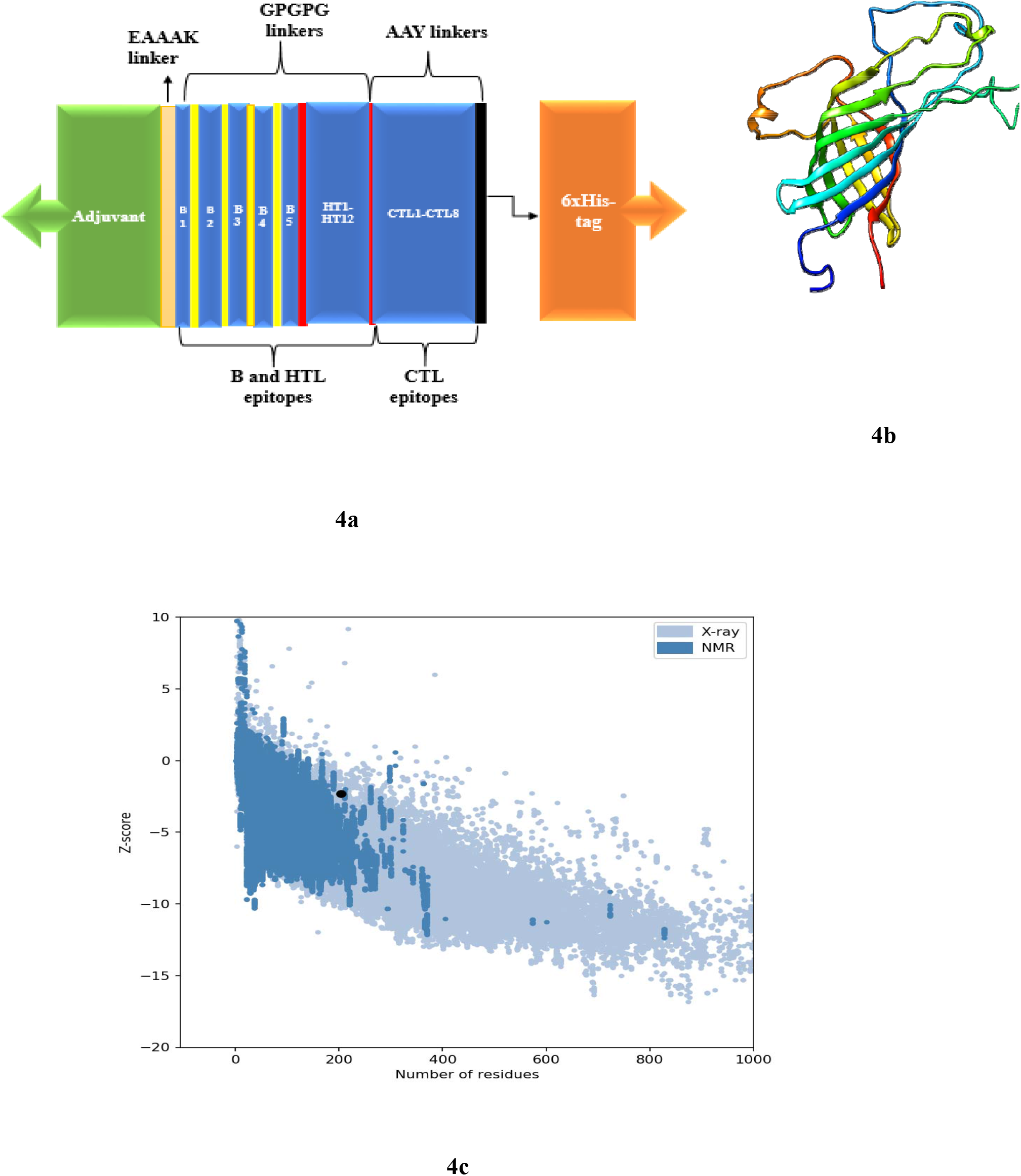
Schematic presentation of the vaccine containing an adjuvant (green) linked with the multi-epitope sequence through an EAAAK linker. The B and HTL epitopes are linked together via the GPGPG linkers while the CTL epitopes are linked with the help of AAY linkers. The 6x-His tag at the carboxyl end. **4b**: Tertiary structure of the vaccine. **4c**: Validation of the structure with a Z score of −2.32. **4d**: Intrinsic solubility profile. Residues lesser than −1 depicts the hydrophobic core of the vaccine peptide.

### Physiochemical, Solubility and Solvation properties of the Vaccine

The physicochemical parameters and solubility properties of a vaccine candidates help to define the efficacy and effectiveness of the vaccine. The molecular weight of the vaccine was 60728.51 Da and the bio-computed theoretical pI was 9.30, with an estimated half-life of 30 hours. The instability index was 27.84, signifying that the vaccine is stable in a solvent environment (>40 signifies instability). The aliphatic index is computed to be 88.21, with a GRAVY score of - 0.056. The intrinsic vaccine solubility at a neutral pH 7 revealed the hydrophilic and hydrophobic core of the vaccine construct **[Figure 4d]**. The overall solubility value of the vaccine was −2.632908 signifying hydrophilic property. The stability of the vaccine construct was assessed considering the radius of gyration and solvent density. The solvent density of the vaccine is 334 e/nm^-3^, the envelope distance is 7 Å, number of q values is 101 and the heavy atoms is a total of 1589. The protein contains 3143 solutes atoms and 14365 water molecules.

The solute has zero charges with the distance of envelope from the solute at 0.7nm while the maximum diameter of the solute is 7.3673 nm **[Figure 4e]**.

**Figure 4e:**
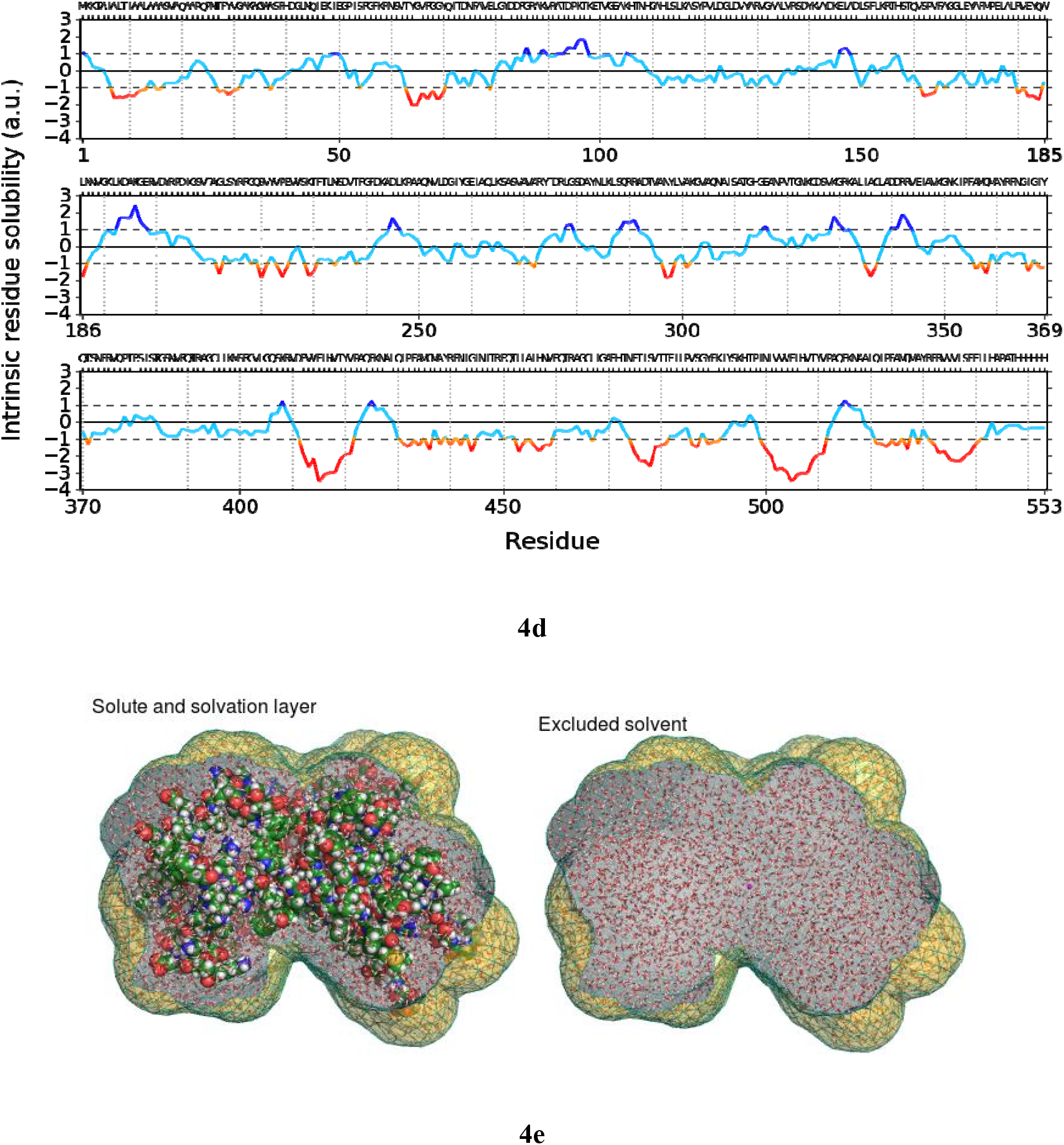
The solute and solvation layer of the vaccine

### Molecular Docking between the Adjuvant Linker of the vaccine and the Toll-like receptor (TLR5)

TLR5 was selected due to its immunomodulatory ability to trigger IFN-β as well as activation of type I IFN responses. This was clearly attested as our selected CD4+ epitopes elicited the Th1 and Th2 cytokines. The molecular interaction between the vaccine and the TLR5 (PDB: 3J0A), was evaluated considering their refined binding energies and various interacting residues. The conformational triggering of the TLR5 receptor was influenced by the adjuvant linker and not the conjugated epitopes. The adjuvant linker binds to the A chain monomer of the toll-like receptor. The interface amino residues of the receptor were: PRO 20, GLN 21, VAL 22, LEU 23, ASN 24, THR 25, PRO 45, and PHE 46 respectively from the A chain, forming a hydrogen bond with the interacting adjuvant residues: ASN 91, GLN 143, HIS 89, ALA 117, LEU 118, VAL 119, ARG 120, THR 142 and SER 141 **[Figure 5]**. The binding energy (ΔG) and dissociation constant (K_d_) predicted values of the protein-protein complex were −12.2 kcal mol^-1^ and 1.0E-09 at 25.0°C respectively.

**Figure 5:**
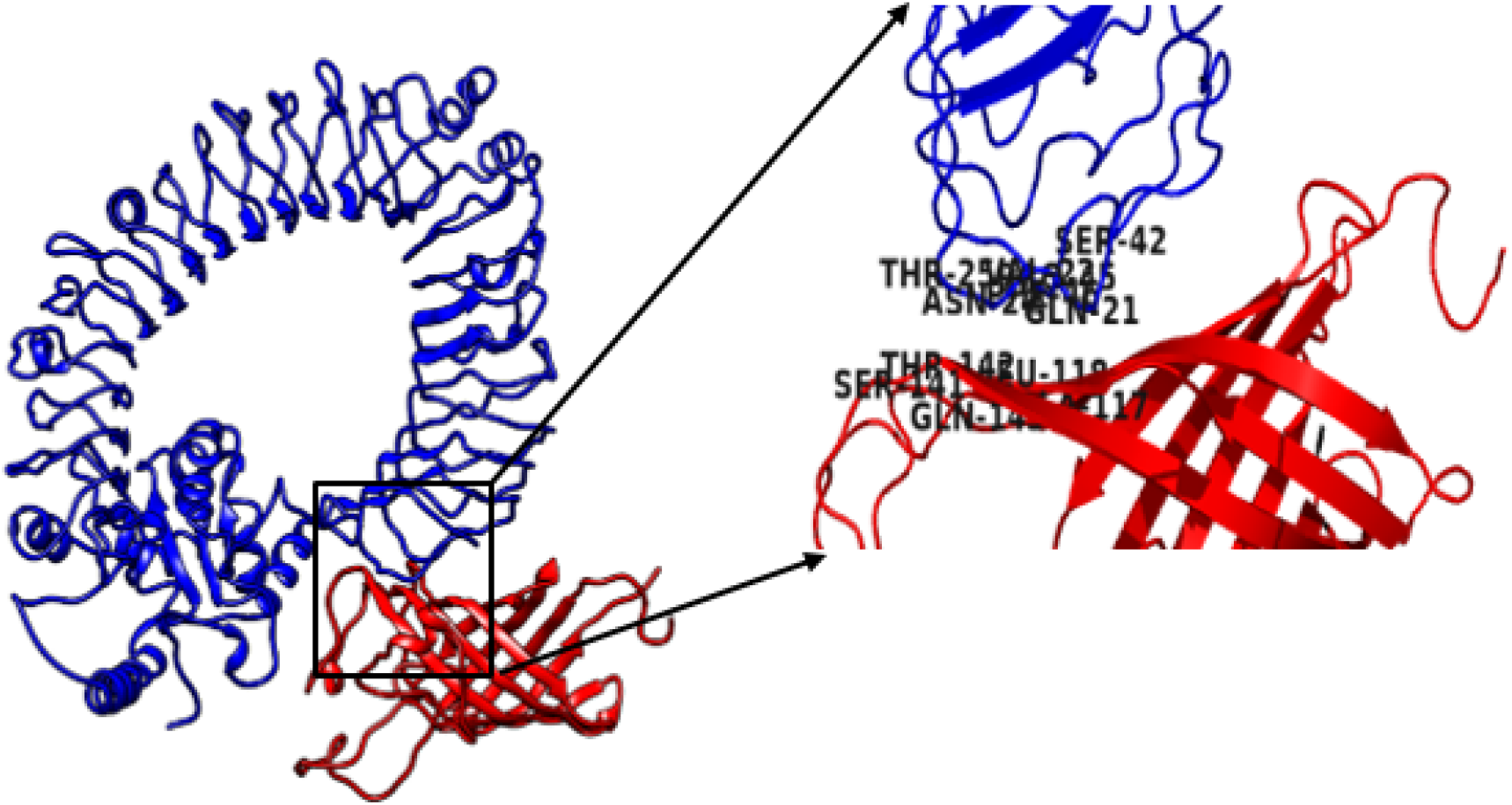
The molecular interaction of the vaccine and TLR5 receptor. The vaccine chain is highlighted in red and the toll like receptor in blue.

### Molecular dynamics simulations

The rigidity of the peptide vaccine system was examined by evaluating the radius of gyration (Rg) values. The analyzed data shows that the average Rg value was 21.0067 nm, indicating that the protein system retained its stability throughout the 85.5 ns time span of the MD simulation **[Figure 6a]**, as this signifies that the peptide vaccine is relatively stable. The molecular interaction between the vaccine peptide and the TLR5 was screened for their protein stability, B factor mobility, and deformity. This analysis relies on the associated coordinates of the docked protein complex **[Figure 6b-6f]**. The eigenvalue found for the complex was 5.812952e-06. The low eigenvalue for the complex signifies easier deformation of the complex, indicating that the docking analysis between the vaccine and the TLR5 will activate immune cascades for destroying the antigens.

**Figure 6a:**
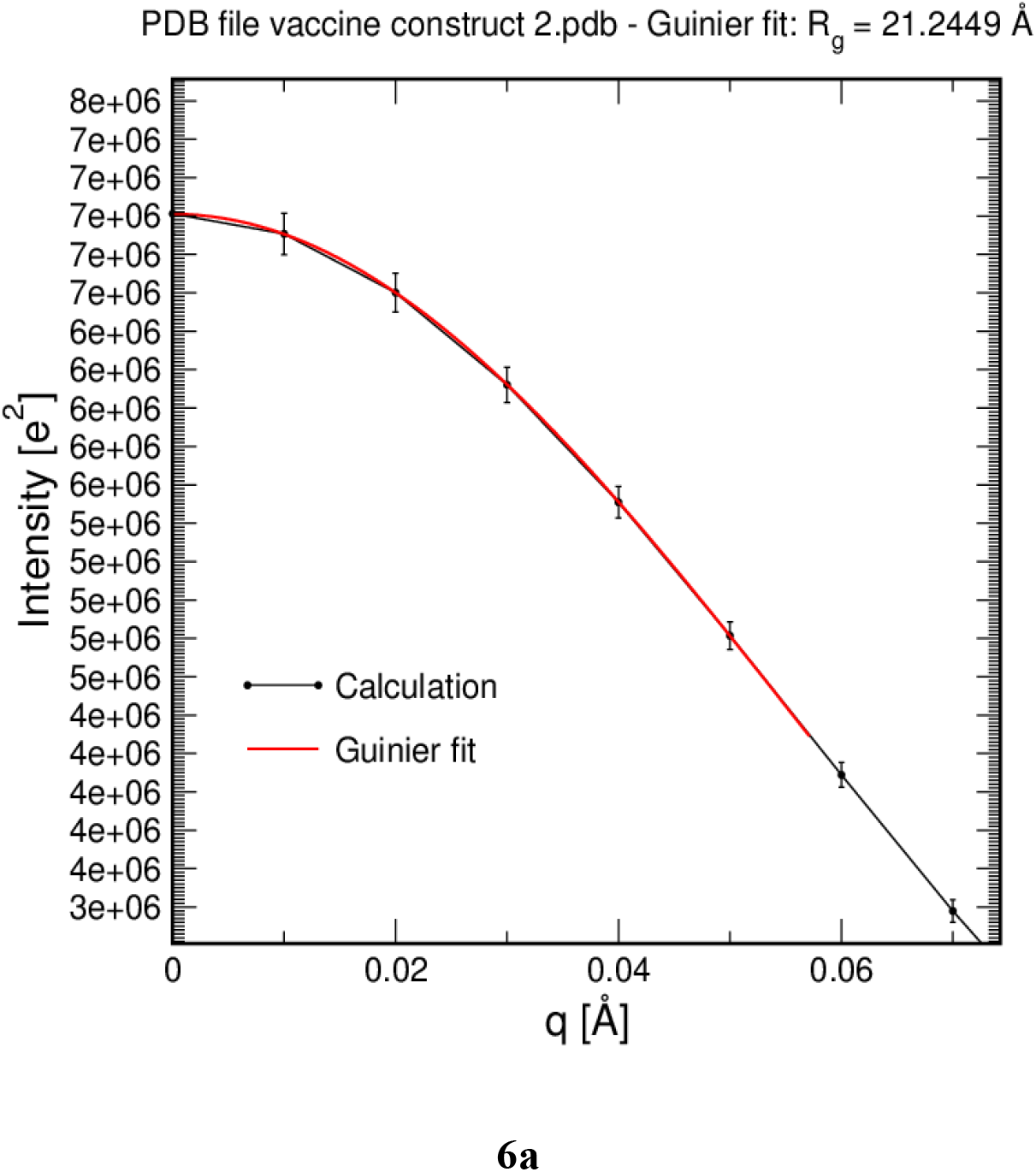
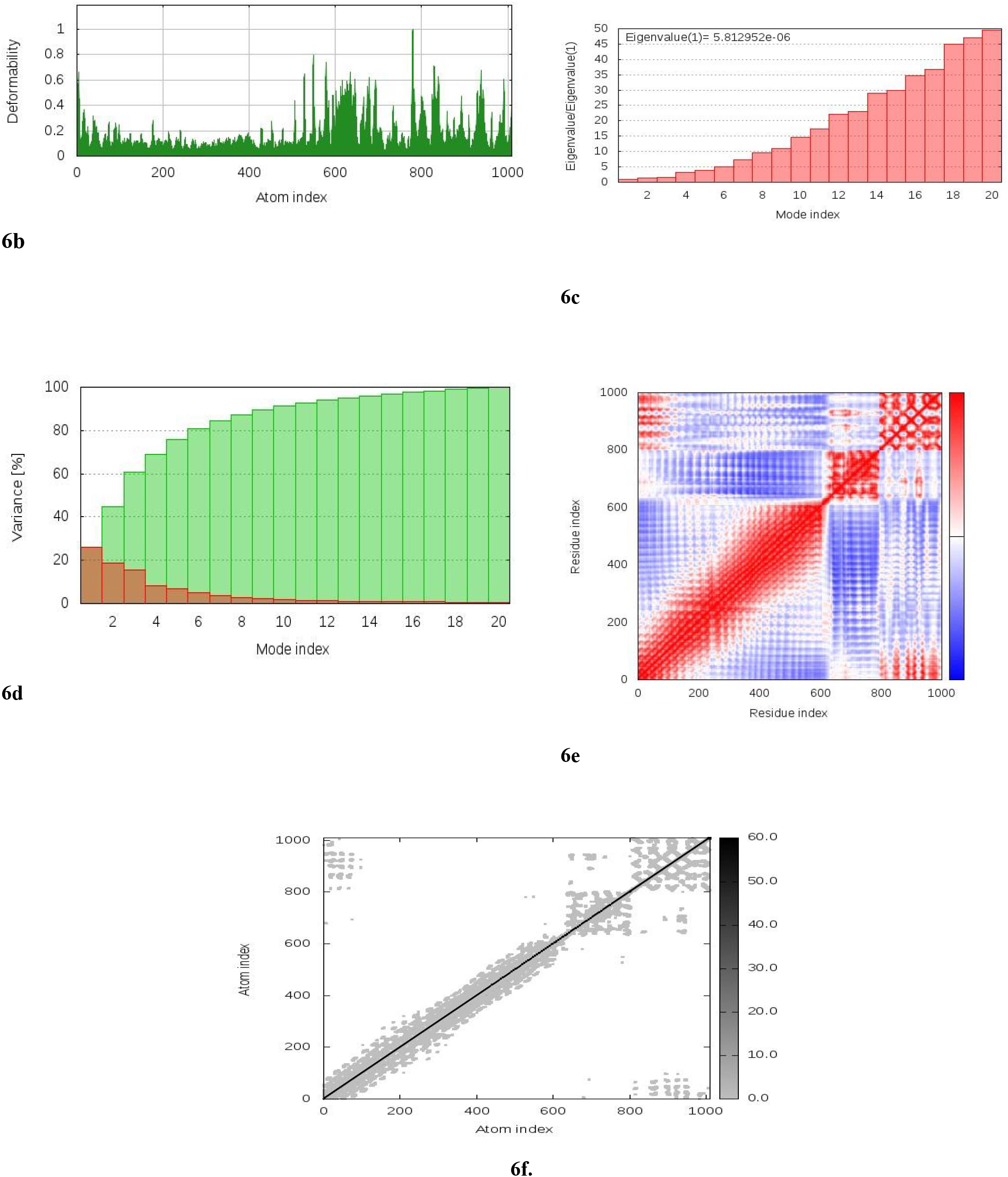
The radius of gyration of the peptide vaccine. **6b-6f**: Molecular dynamics simulation of the vaccine-TLR5 complex, showing (a) eigenvalue; (b) deformability; (c) B-factor; (d) Covariance matrix indicates coupling between pairs of residues (red), uncorrelated (white) or anti-correlated (blue) motions. and (e) elastic network analysis which defines which pairs of atoms are connected by springs.

### Codon Optimization and *In Silico* Cloning

The length of the optimized vaccine codon sequence was 1659 nucleotides. The GC content of the cDNA sequence and codon adaptive index was calculated as 50.8%, which still falls within the recommended range of 30-70%, for effective translational efficiency. The codon adaptive index was calculated as 0.93, falling within the range of 0.8-1.0, signifying the effective expression of the vaccine constructs in the *E. coli*. EagI-NotI and SAlI sites were subsequently cloned into the pET28a (+) vector. The estimated length of the clone was 7.028 kbp [**Figure 7**].

**Figure 7:**
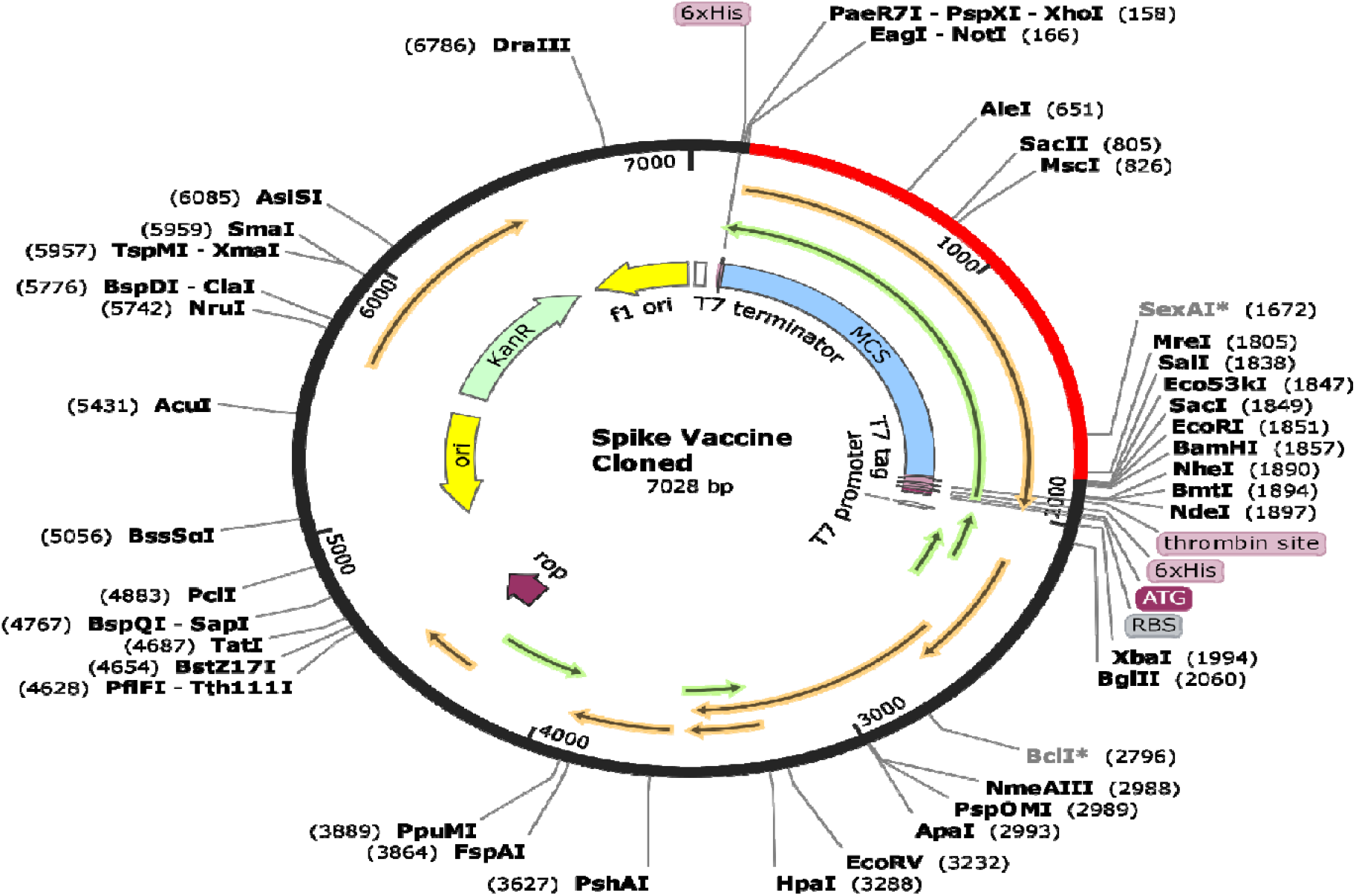
*In silico* cloning of the final vaccine construct into pET28a (+) expression vector where the red part indicates the coding gene for the vaccine surrounded between EagI-NotI (166) and SAlI (1838) while the vector backbone has shown in a black circle. MCS represents the multiple cloning site.

### Immune Simulation of the Chimeric Peptide Vaccine

At every administration and repetitive exposure to the attenuated peptide vaccine, there was a significant increase in the antibody response with a simultaneous decrease in the antigen level. This is characterized by an increase in IgM concentration. The observed increase in immunoglobin activities involving the combination of IgG1 + IgG2, IgG + IgM and IgM antibodies components, was another evidence that the vaccine stimulated a good immune response **[Figure 8a]**. This same immune response was observed for the B cell population, with increase in the population of B cell memory formation. The subsequent increase in cytokine levels was also induced by the vaccine [**Figure 8b**]. Similar elevated responses were observed for the CD4^+^ and CD8^+^ cells population with a significant increase in memory formation **[Figure 8c-8f]**. The concomitant increase in dendritic cells and natural killer cells were another good immune response attribute exhibited by the vaccine **[Figure 8g-8j]**.

**Figure 8a:**
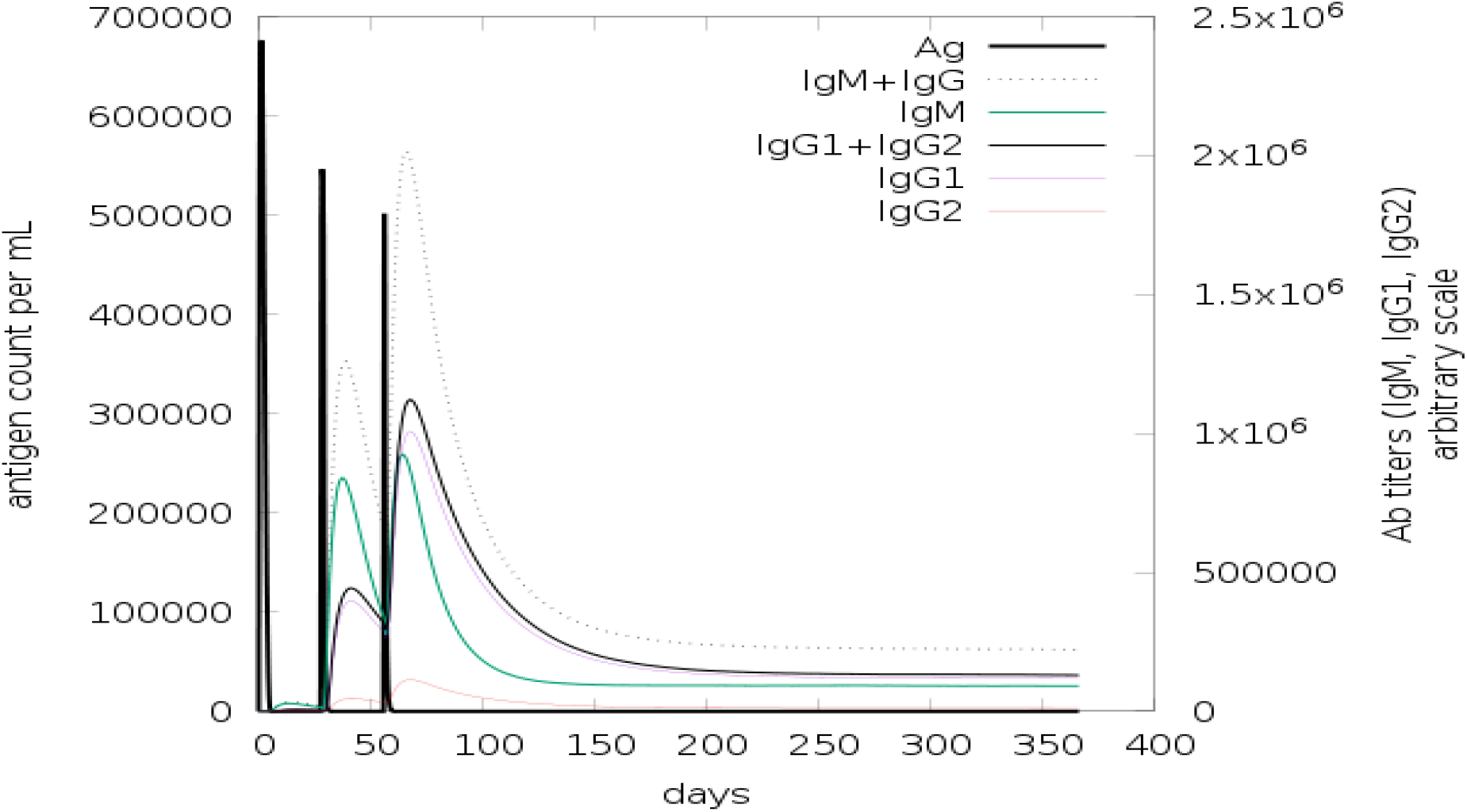
Increase in immunoglobin activities and concomitant decrease in antigen level in 365 days of vaccination. Antibodies are sub-divided per isotype.

**Figure 8b:**
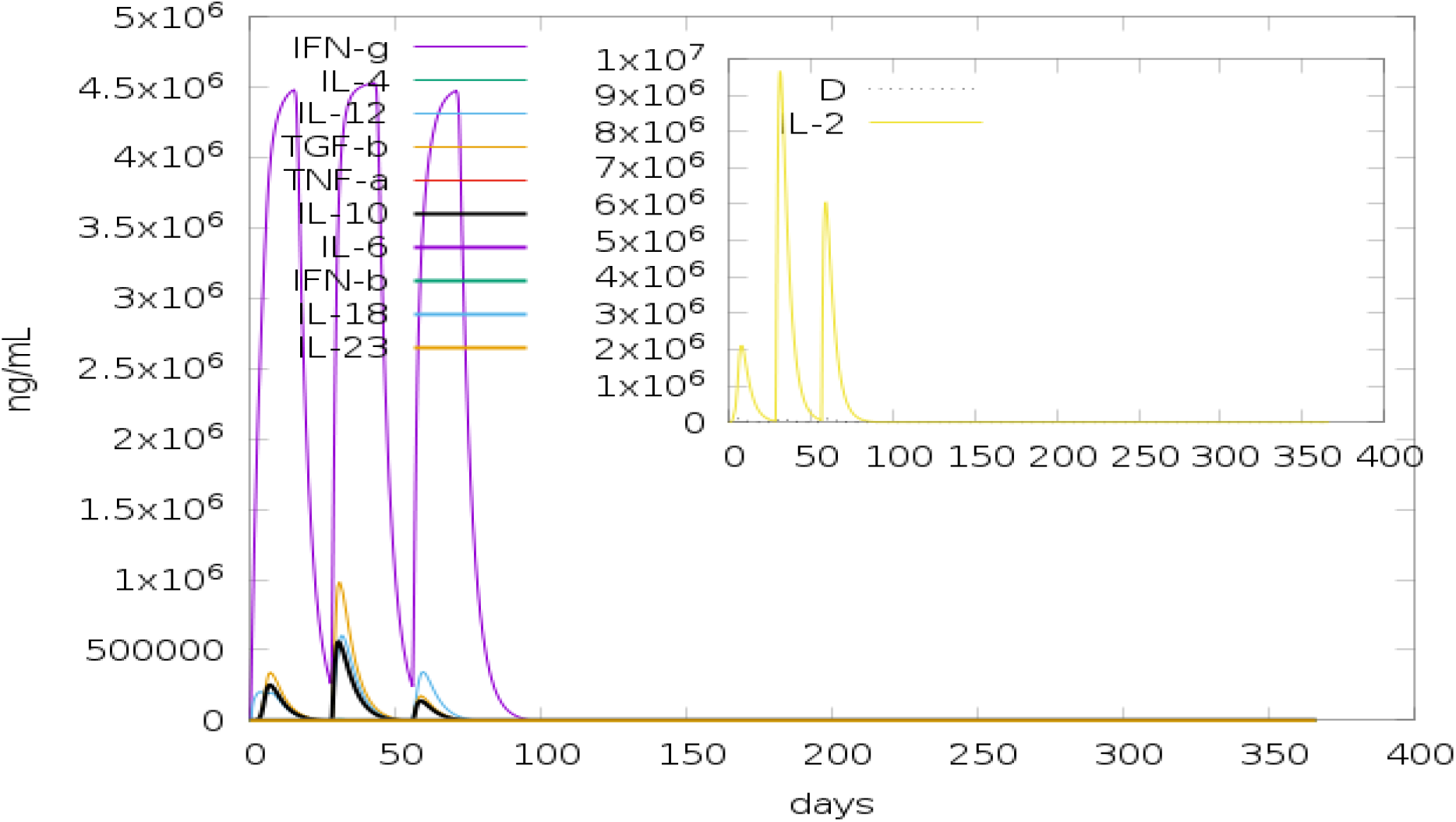
Concentration of cytokines and interleukins. D in the inset plot is danger signal

**Figure 8c-8i:**
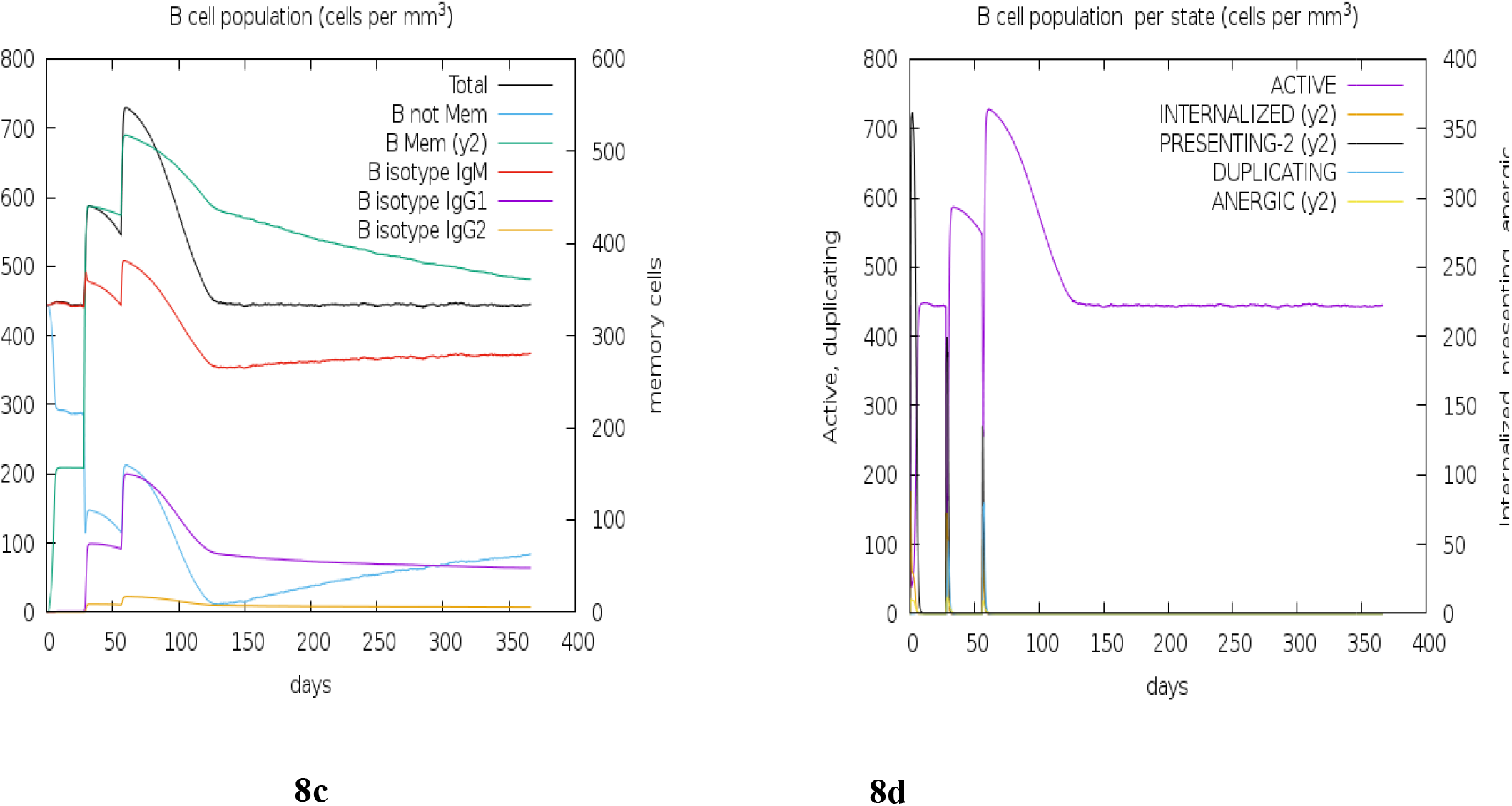
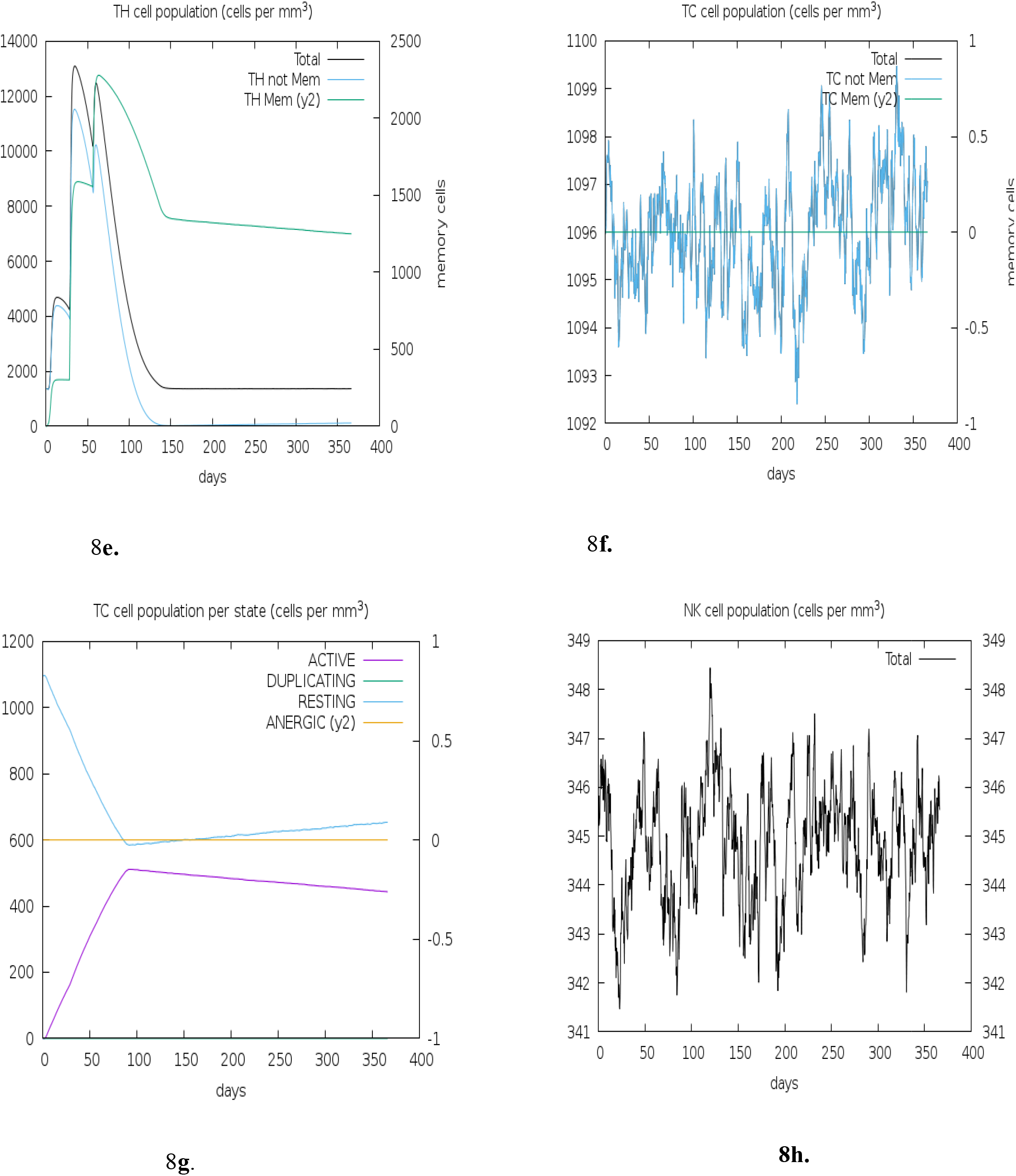
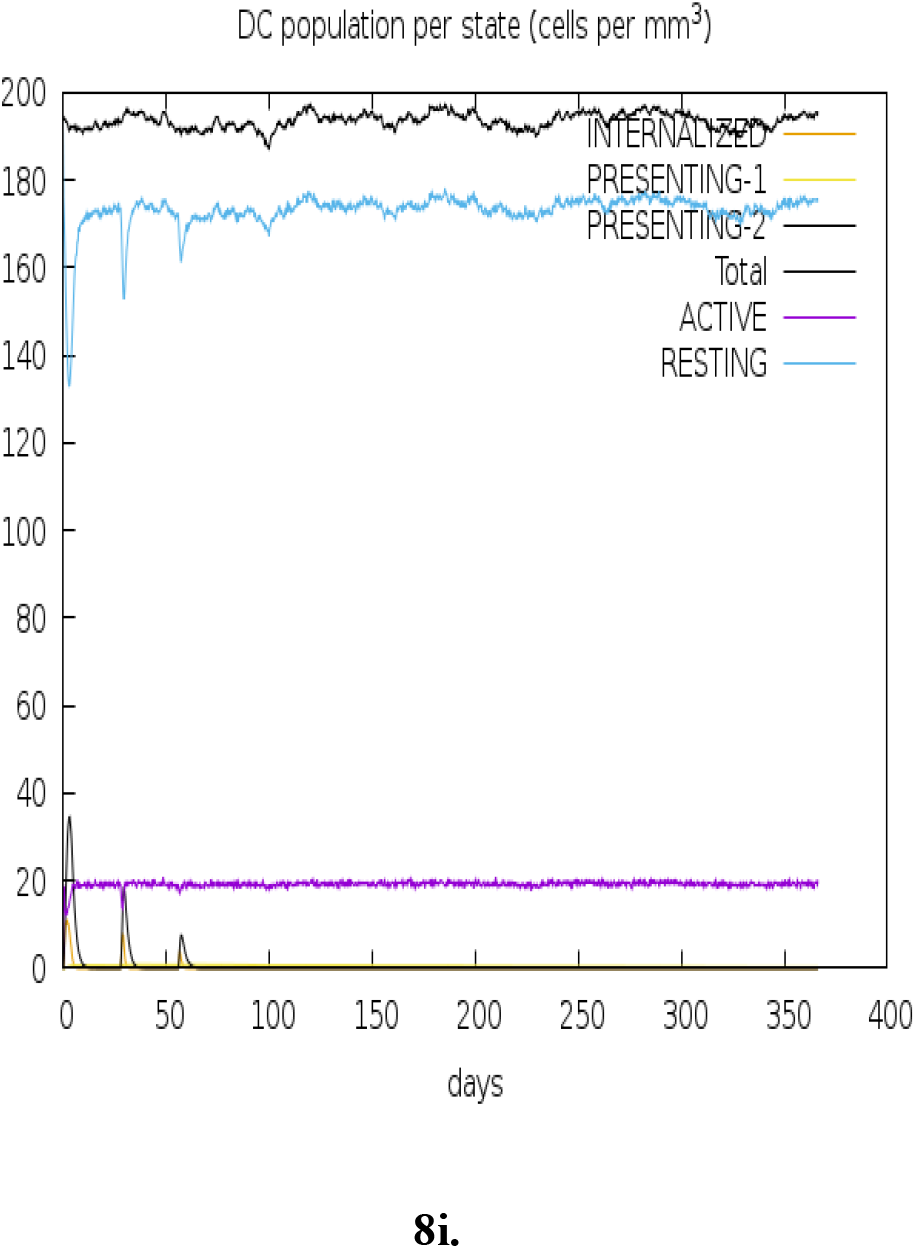
**c.** B lymphocytes: total count, memory cells, and sub-divided in isotypes IgM, IgG1 and IgG2. **d.** CD4 T-helper lymphocytes count. **e.** CD4 T-helper lymphocytes count. **f.** CD8 T-cytotoxic lymphocytes count. **g.** CD8 T-cytotoxic lymphocytes count per entity-state. **h.** Natural Killer cells (total count). **i.** Dendritic cells.

## DISCUSSION

The emergence of new coronavirus strain SARS-CoV-2 viral diseases is a global threat, responsible for the death of many across the globe, including health care workers **[1]**. Therefore, there is an urgent need for therapeutics and preventive measures that could confer protection against this enigma. Our study was therefore centered on using an epitope peptide-based vaccine design against the SARS-CoV-2 spike protein complex. We successfully developed a peptide vaccine after a rigorous round of screenings and conditions in selecting the epitopes, using an array of immune-informatics tools. Criteria such as the elicitation of immune response with lesser or no potential infectious abilities are taken into consideration before each epitope selection, using stipulated thresholds. Both the B cell and T cell epitopes were predicted in this study. B cells recognize solvent-exposed antigens through antigen receptors, named as B cell receptors (BCR), consisting of membrane-bound immunoglobulins **[38]**. The B cell epitopes were selected based on surface accessibility, and Kolaskar and Tongaonkar antigenicity scale methods. Five antigenic B cell epitopes were predicted in the study. The peptide also has a nontoxicity attribute, making it a safe vaccine candidate.

The T cell antigenic epitopes capable of binding a large number of MHC I and MHC II alleles were predicted using various tools, and selections were made based on the recommended thresholds. T cell epitopes presented by MHC class I molecules are typically peptides between 8 and 11 amino acids in length, whereas MHC class II molecules present longer peptides, 13-17 amino acids in length [**39**]. The CD8^+^ T cell recognizes the antigen of a pathogen after its attachment with the MHC I molecules, therefore triggering a cytotoxic response against the pathogen [**40**]. Eight promising CD8^+^ T cell epitopes were predicted. These peptides are capable of eliciting a cytotoxic response with their respective antigenic properties.

Twelve helper T cell epitopes were predicted based on their virtual attachments with the beta chain of antigen-presenting major histocompatibility complex class II (MHC II) molecule. In complex with the beta chain HLA-DR, the T cell epitopes display antigenic peptides on professional antigen-presenting cells (APCs) for recognition by alpha-beta T cell receptor (TCR) on HLA-DRB1-restricted CD4-positive T cells [**40, 41**]. This guides antigen-specific T helper effector functions, both antibody-mediated immune response and macrophage activation, to ultimately eliminate the infectious agents and transformed cells [**40**]. The 12 selected epitopes were highly antigenic, with each capable of inducing any of the Th1 and Th2 cytokines, which corroborates the findings that T cell CD4+ is capable of inducing an adaptive immune response in the human cells **[41]**, as adaptive immunity is articulated by lymphocytes, more specifically by B- and T-cells, which are responsible for the humoral and cell-mediated immunity [**41**]. Prediction of peptide binding to MHC II molecules readily discriminate CD4 T-cell epitopes, but cannot tell their ability to activate the response of specific CD4 T-cell subsets (e.g., Th1, Th2, and Treg). However, there is evidence that some CD4 T-cell epitopes appear to stimulate specific subsets of Th cells **[40, 41]**. The ability to distinguish the epitopes capable of inducing distinct responses is highly imperative in vaccine development. This study was able to differentiate the abilities of each MHC class II epitopes of eliciting either the Th1 and Th2 cytokines or both.

The docking scores involving the predicted epitopes and the MHC II molecules were comparatively evaluated. Considering the docking attributes of the MHC II epitopes, they displayed a varied binding putative attribute. Some of the epitopes core peptides had a flanking region away from the MHC class II binding groove. Generally, MHC-II peptides contain a central binding motif of nine core amino residues that specifically attach to the MHC II binding groove. These core peptides interact with the allelic specific pockets of the MHCII binding groove.

Considering the population cover for the MHC-I and II epitopes, the European population significantly show a potential response to the selected epitopes, however, we suggest that the IEDB population coverage tool have less MHC class I and II deposition of alleles from continents like Africa and Asia, compared to the European and Americans. Immunization of the MHCII T cell epitopes will confer protection to 80.88% of the world population, while MHCII T cell epitopes will confer protection to 55.23%. It is imperative to know that specific interactions with high binding affinity epitope / HLA allele class II molecule unleash protective and specific adaptive immune response [**39–42**].

The designed vaccine construct was predicted to be stable, soluble (i.e. hydrophilic) and with increased thermostability, as depicted in its physicochemical characteristics. The molecular weight of the vaccine and its high pI value signifies the efficacy as well as the stability of the vaccine construct since proteins having <110kD molecular weight are considered good vaccine candidates [**43, 44**]. Apart from size, surface properties like surface charge and hydrophobicity can affect a designed vaccine candidate. Neutral or negatively charged molecules are preferred and a balance between its hydrophobicity and hydrophilicity is crucial in designing vaccine candidates [**45**].

### Conclusion

This is a novel approach to predicting SARS-CoV-2 epitope peptide-based vaccine targeting the spike protein, utilizing immune-informatics tools and immune simulation measures. These predicted antigenic epitopes would hasten the production of protective vaccine for patients around the world whose immune system has been compromised. Our selected epitopes (B and T cells) will constitute a good vaccine candidate against the spike protein. In future studies, other effective stimulants that could aid the rapid response of cells to antigens will be considered and assessed.

## Acknowledgement

We sincerely appreciate the effort of Sblend Digital overseen by Mr. Agboola Olamide, for providing some of the sophisticated computing accessories and work space to conduct this research.

## Data Availability

Data are available upon request and may be obtained by contacting the corresponding author

## Supplementary Data

**Figure S1a-S1c:**
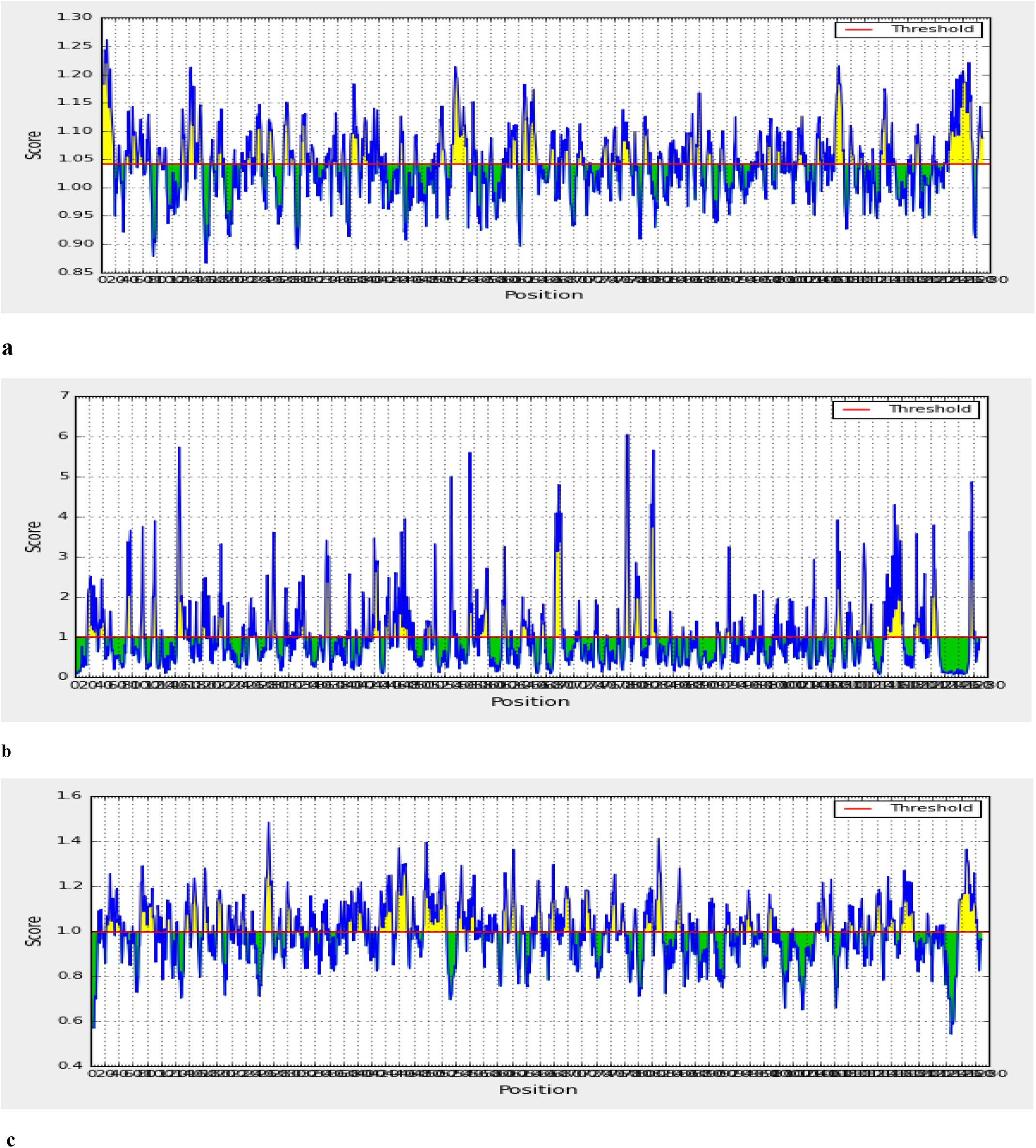
Prediction of antigenic determinants. **a.** Kolaskar and Tongaonkar antigenicity scale. **b.** Emini surface accessibility **c.** Chou and Fasman beta turns. Green regions under the threshold color denotes unfavourable related to the properties of interest. Yellow colours are above the threshold sharing higher scores. Horizontal red lines represent the threshold.

### Physiochemical analysis of the SARS-CoV-2 Spike Glycoprotein

The sequence of the primary structure of SARS-CoV-2 spike glycoprotein was computed, analyzed and tabulated **[Table 1].** The molecular weight was estimated at 141178.47 Da. To calculate the extinction coefficient, wavelengths of varying amount (276, 278, 279, 280 and 282 nm) were computed. But wavelength at 280nm is usually favored because of high protein absorption. So, the extinction coefficient at 280nm is 148960 M^-1^cm^-1^ with respect to the cysteine, trypsin and tyrosine concentrations. The spike glycoprotein is highly stable as the instability index was 33.01, because protein instability index at or above 40 is considered not stable. The isoelectric point or computed theoretical pI of the spike glycoprotein was acidic, lower than 7. The information on the theoretical pI is useful in developing buffer system for the purification of recombinant protein. The total number of negatively charged residues (Asp + Glu) is 110, while that of positively charged residues (Arg + Lys) is 103. The positively charged residues are lesser than the negatively charged counterparts which signifies that the protein is intracellular. The half-life of the protein is 30hours, while the aliphatic index which is the relative volume occupied by aliphatic side chains such as valine, isoleucine, alanine and leucine, is 84.67. At such value, the protein has a high thermostability. The Grand Average hydropathy (GRAVY) of a protein is calculated as the sum of hydropathy values of all amino acids, divided by the number of residues in the sequence. The gravy value of the spike glycoprotein is −0.079, depicting its hydrophilic nature, and better interaction with water. The individual amino composition of the protein is summarized [**Figure 1**]. Every individual amino residue plays a role in the protein function, structure and signaling, depending on their position. The four major amino residues were leucine, serine, threonine and valine. Serine and threonine majorly perform the phosphorylation function which is expedient for the protein’s signaling pathway, as they have hydroxyl functional group with affinity for phosphate group. The least amino residues were Trp (0.90%), Met (1.10%) and His (1.3%).

**S-Table 1:**
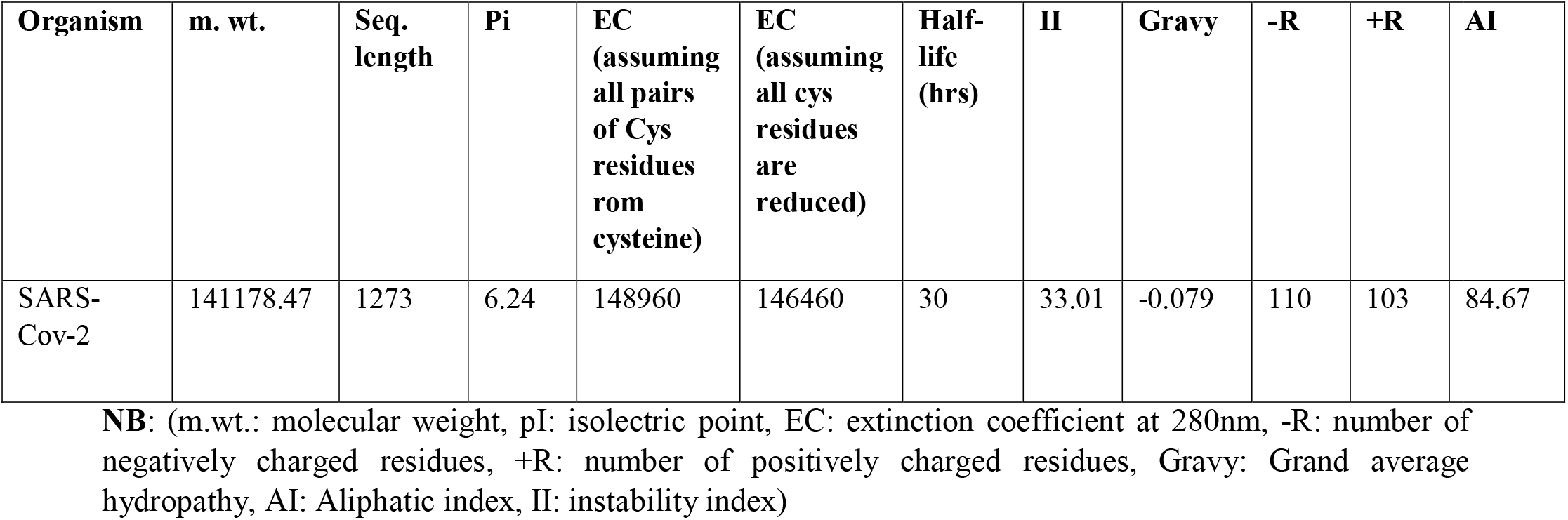
Physicochemical properties of SARS-CoV-2 Spike glycoprotein

**Figure S2:**
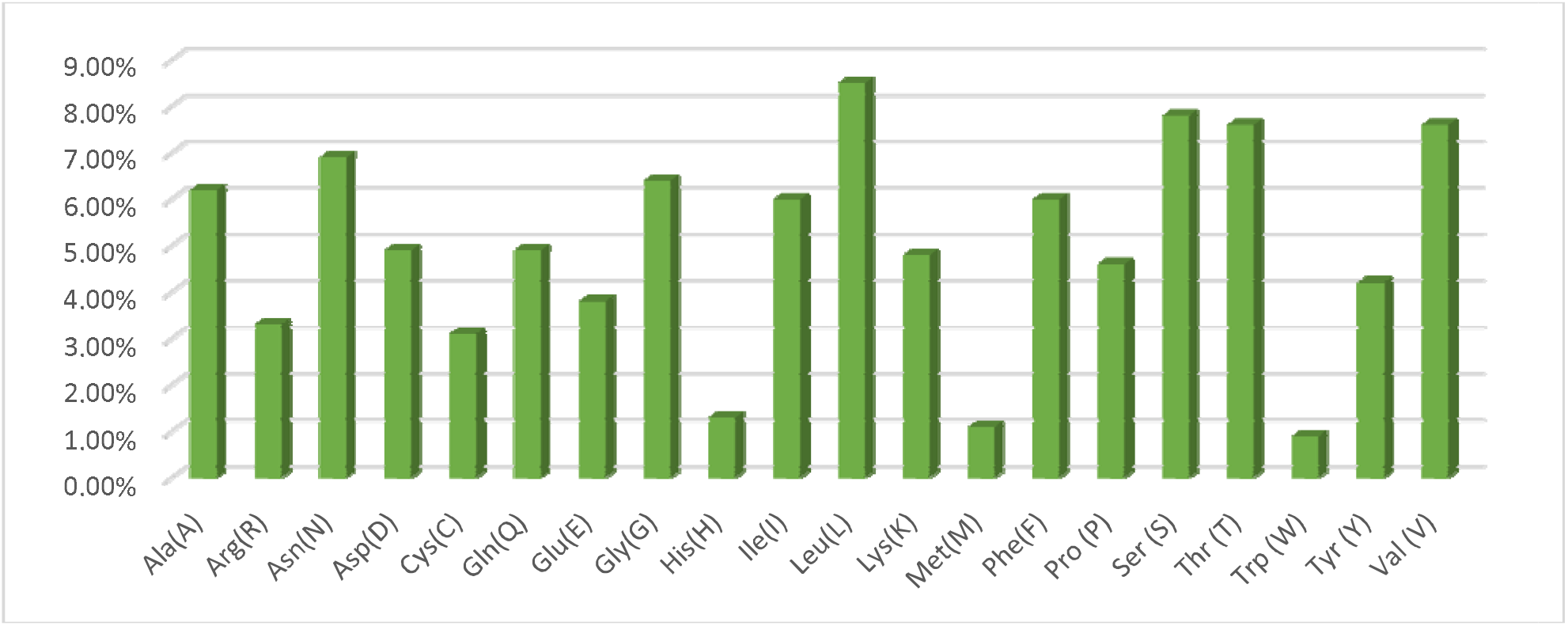
Percentage of amino acids present in SARS-CoV-2 Spike glycoprotein.

**S-Table 2:** Discontinous B cell epitope contact numbers in the spike glycoprotein.

| Chain ID | Residue ID | Residue Name | Contact Number | Propensity Score | Discotope score |
| --- | --- | --- | --- | --- | --- |
| A | 809 | PRO | 8 | 2.079 | -2.909 |
| A | 810 | SER | 7 | 2.066 | -2.786 |
| A | 794 | ILE | 7 | 2.211 | -2.745 |
| A | 793 | PRO | 7 | 2.483 | -2.727 |
| A | 560 | LEU | 7 | 0.591 | -2.39 |
| A | 1145 | LEU | 7 | 0.714 | -1.921 |
| A | 491 | PRO | 6 | 0.255 | -1.434 |
| A | 562 | PHE | 7 | 0.773 | -1.289 |
| A | 1146 | ASP | 6 | 0.61 | -1.017 |
| B | 1144 | GLU | 8 | 0.918 | -3.082 |
| B | 811 | LYS | 9 | 1.491 | -3.009 |
| B | 560 | LEU | 7 | 0.591 | -2.909 |
| B | 1145 | LEU | 7 | 0.714 | -2.786 |
| B | 1146 | ASP | 6 | 0.61 | -2.39 |
| B | 809 | PRO | 8 | 2.079 | -1.921 |
| B | 810 | SER | 7 | 2.066 | -1.434 |
| B | 794 | ILE | 7 | 2.211 | -1.289 |
| B | 793 | PRO | 7 | 2.483 | -1.017 |
| C | 792 | PRO | 10 | 1.669 | -3.331 |
| C | 1144 | GLU | 8 | 0.918 | -3.082 |
| C | 811 | LYS | 9 | 1.491 | -3.009 |
| C | 1145 | LEU | 7 | 0.714 | -2.786 |
| C | 1146 | ASP | 6 | 0.61 | -2.39 |
| C | 810 | SER | 8 | 1.996 | -2.004 |
| C | 809 | PRO | 8 | 2.079 | -1.921 |
| C | 794 | ILE | 7 | 2.211 | -1.289 |
| C | 793 | PRO | 7 | 2.483 | -1.017 |

Residues are shown in three-letter code, and number of contacts shows the connection of amino acid with neighbouring groups

**Figure S3:**
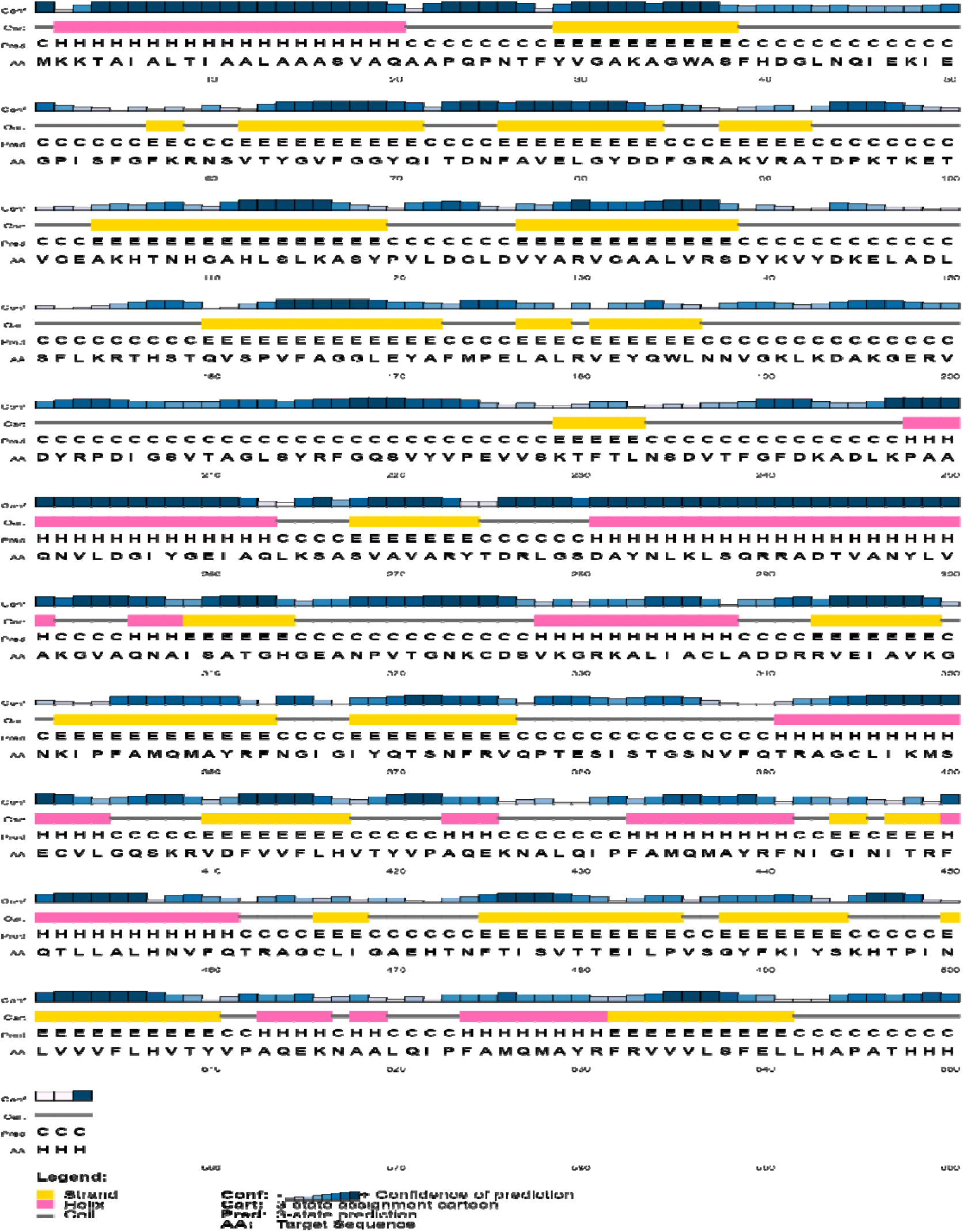
Diagrammatic representation of secondary structure prediction of the vaccine construct. Here, the β-strands, α-helix and random coils are indicated by yellow, pink and blue colour, respectively.

**Figure S4:**
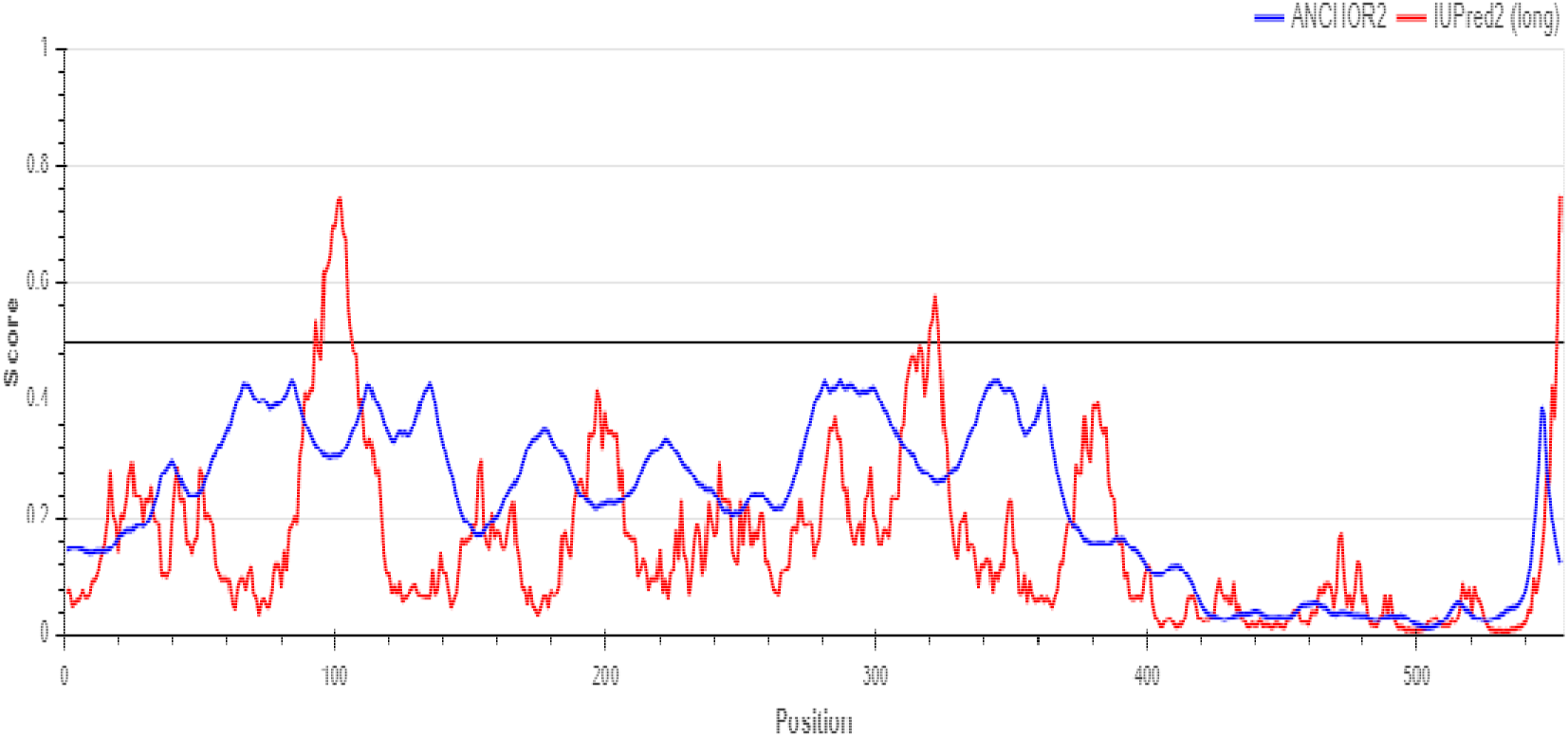
Disordered region of the peptide vaccine construct. Regions (lines) exceeding the threshold of 0.5, was considered disordered.

